# Deep Learning Analysis on Images of iPSC-derived Motor Neurons Carrying fALS-genetics Reveals Disease-Relevant Phenotypes

**DOI:** 10.1101/2024.01.04.574270

**Authors:** Rahul Atmaramani, Tommaso Dreossi, Kevin Ford, Lin Gan, Jana Mitchell, Shengjiang Tu, Jeevaa Velayutham, Haoyang Zeng, Michael Chickering, Tom Soare, Srinivasan Sivanandan, Ryan Conrad, Yujia Bao, Santiago Akle, Jonathan Liu, Stephanie Redmond, Syuan-Ming Guo, Patrick Conrad, Flora Yi, Nick Atkeson, Difei Xu, Aidan McMorrow, Emiliano Hergenreder, Mukund Hari, Ahmed Sandakli, Nitya Mittal, Liyuan Zhang, Aaron Topol, Brigham Hartley, Elaine Lam, Eva-Maria Krauel, Theofanis Karaletsos, Mark Labow, Richard Hargreaves, Matthew Trotter, Shameek Biswas, Angela Oliveira Pisco, Ajamete Kaykas, Daphne Koller, Samuel Sances

## Abstract

Amyotrophic lateral sclerosis (ALS) is a devastating condition with very limited treatment options. It is a heterogeneous disease with complex genetics and unclear etiology, making the discovery of disease-modifying interventions very challenging. To discover novel mechanisms underlying ALS, we leverage a unique platform that combines isogenic, induced pluripotent stem cell (iPSC)-derived models of disease-causing mutations with rich phenotyping via high-content imaging and deep learning models. We introduced eight mutations that cause familial ALS (fALS) into multiple donor iPSC lines, and differentiated them into motor neurons to create multiple isogenic pairs of healthy (wild-type) and sick (mutant) motor neurons. We collected extensive high-content imaging data and used machine learning (ML) to process the images, segment the cells, and learn phenotypes. Self-supervised ML was used to create a concise embedding that captured significant, ALS-relevant biological information in these images. We demonstrate that ML models trained on core cell morphology alone can accurately predict TDP-43 mislocalization, a known phenotypic feature related to ALS. In addition, we were able to impute RNA expression from these image embeddings, in a way that elucidates molecular differences between mutants and wild-type cells. Finally, predictors leveraging these embeddings are able to distinguish between mutant and wild-type both within and across donors, defining cellular, ML-derived disease models for diverse fALS mutations. These disease models are the foundation for a novel screening approach to discover disease-modifying targets for familial ALS.

## Introduction

Cellular models of disease, while intrinsically reductionist, have provided multiple deep insights on disease processes, and continue to enable at-scale screening for disease-modifying treatments. Generating a good disease model requires that we build a robust cellular system that encompasses relevant biology, introduce perturbations that reflect causal disease processes, and measure and identify the phenotypic consequences of the disease. Each of these steps can be quite challenging: biologically relevant cellular systems are often hard to generate, leading to the use of less translatable systems such as cancer cell lines; disease etiology is not always well-understood and in many cases is heterogeneous; and disease phenotypes can be multifactorial and subtle. In this paper, we present a novel platform that helps address these challenges for diseases with a strong genetic basis. Our platform integrates a set of cutting-edge technologies: robust induced pluripotent stem cell (iPSC)-derived models to generate biologically-relevant cellular systems; genome editing to introduce disease-causing mutations into multiple lines to create well-powered isogenic “case-controls”; and machine learning (ML) to identify subtle, yet reproducible, disease-associated phenotypes in high-content imaging (HCI) data.

We deploy this platform towards the interrogation of amyotrophic lateral sclerosis (ALS), a devastating neurodegenerative disease with no disease-modifying treatments. Clinical presentation and progression of ALS in patients is highly variable, and is driven by multiple genetic contributors spanning diverse pathways. At the same time, these diverse causal factors ultimately result in shared clinical manifestations driven by motor neuron (MN) cell death in familial and sporadic ALS (Masrori & Van Damme, 2020). Previous ALS studies have also described shared *in vitro* functional and molecular phenotypes, including TDP-43 mislocalization, which is present in >95% of ALS patient tissues at post mortem examination. However this *in vitro*-derived signal is quite subtle, sensitive to genetic background, and variable in reproducibility across studies from different familial ALS (fALS) patient lines (Bilican et al., 2012). Given the diversity in function of fALS genes, it is also likely that a spectrum of pathogenic mechanisms underlie their contribution to hallmark pathology *in vitro*. But these insights are not visible if we focus only on a few known anchor phenotypes such as human-defined, one-dimensional traits like TDP-43 mislocalization.

### Generating robust, broad-based in vitro disease models for ALS

To more robustly and comprehensively interrogate ALS genetics, we first developed an automated workflow utilizing a rapid (10-day) differentiation protocol that produces functional iPSC-derived motor neurons (iMNs) at high quality and purity. We interrogated a breadth of fALS disease processes by selecting seven known fALS-causing genes, chosen because they have been found both to exhibit strong monogenic signals and to result in a range of cellular processes potentially related to ALS pathogenesis (Brown & Al-Chalabi, 2017; Masrori & Van Damme, 2020; Weishaupt et al., 2016). The range in pathogenic mechanisms were of particular interest, because they create the opportunity to capture potentially separate pathogenic mechanisms related to ALS *in vitro* using ML classification techniques. Specifically, we selected fALS mutations in C9ORF72 and SOD1 because they are found in the highest prevalence in fALS patient populations (Akçimen et al., 2023); C9ORF72, TARDBP, VCP, and TBK1 because they are known to have functions related to TDP-43 (Shao et al., 2022); DCTN1 due to evidence of its role in axonal transport-related deficits (Castellanos-Montiel et al., 2020); and SOD1 and FUS because they do not follow TDP-43 pathology and therefore represent unique clinical presentation (Mackenzie et al., 2010); (**Figure 1.A**). We also included two fALS-causing mutations in the TARDBP gene (G295S and M337V) in our model to help us define common phenotypic effects of these mutations. In most cases, disease-associated mutations were edited into wild-type (WT) iPSCs to produce isogenic case-controls; for C9ORF72, however, we corrected the disease-causing mutations to create the isogenic pairs. These lines were differentiated into lower-motor-neuron-like cells via our automated protocol, forming the basis of the multi-mutant fALS disease model for ML-derived phenotypic discovery.

**Figure 1.**
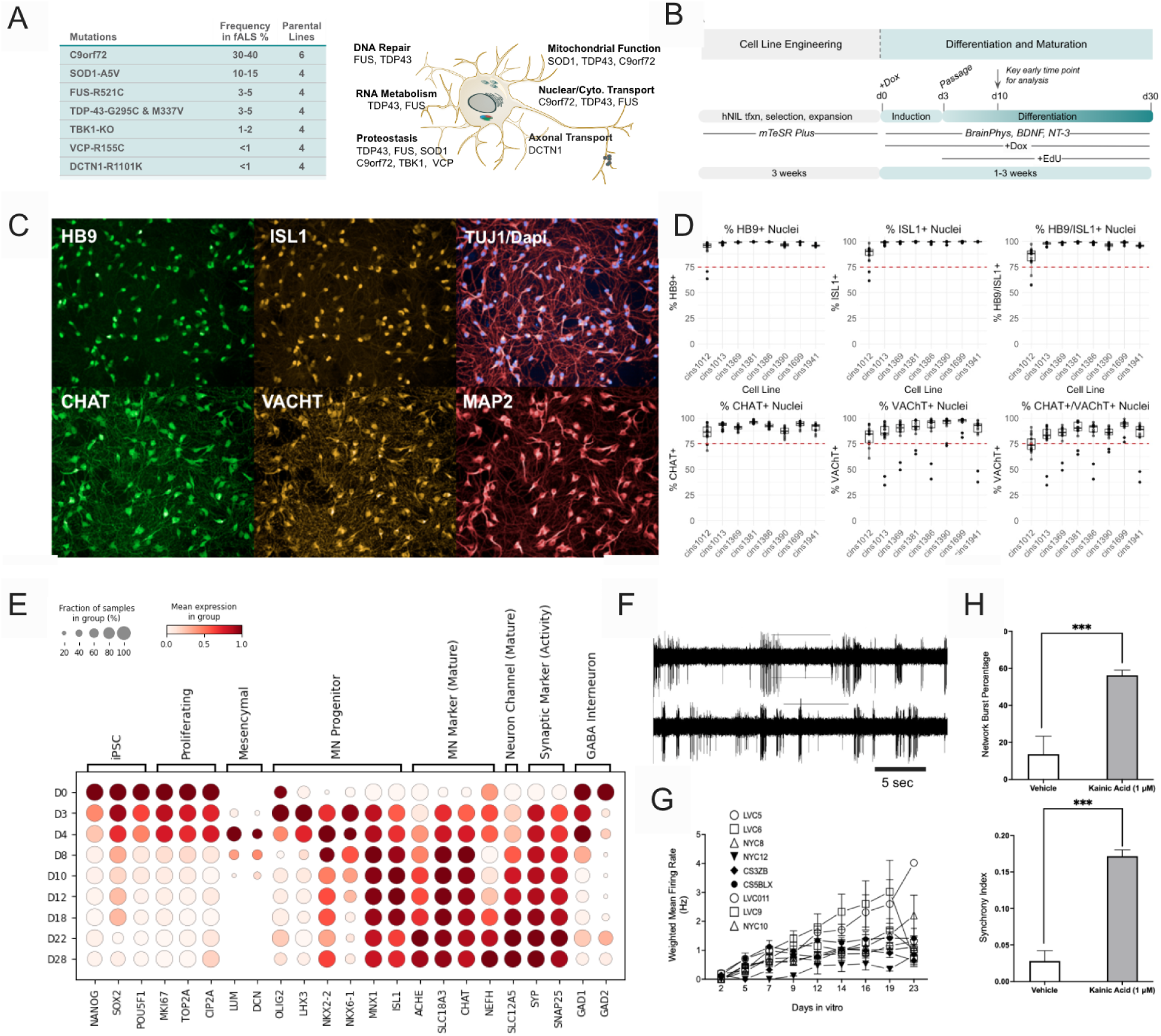
FALS mutants engineered from iPSCs to ALS-relevant hNIL motor neurons. **A.** Eight mutations from seven ALS-causing genes for phenotypic discovery. Genes were selected based on the prevalence of the genes in the ALS population, observed phenotypes in *in vitro* iPSC-MNs, and distinct cellular functions such as RNA metabolism, DNA repair, and proteostasis (right panel). **B.** Schematic of differentiation process for human transcription factor (hNIL)-based expression through piggyBac and doxycycline induction. Dox is doxycycline; EdU is the mitotic inhibitor 5-ethynyl-2-deoxyuridine; BDNF is brain-derived growth factor; NT-3 is neurotrophin-3. mTeSR plus and BrainPhys are cell media (Bardy et al., 2015). **C.** Representative fluorescent images of hNIL neurons cultured to day 10, fixed and stained for relevant motor neuron markers. Nuclear transcription factors Homeobox containing 9 (HB9 (aka MNX1)), islet 1 (ISL1), cytoplasmic beta III tubulin (TUJ1, aka (TUBB3)) and MAP2, and MN functional proteins Choline Acytltransferase (CHAT), and vesicular CHAT (VCHAT). Antibodies are listed in methods. **D.** Quantification of cell masks generated from fluorescent images across individual donor lines (x-axis) reported as a percentage of total DAPI positive nuclei or total cells. Red dashed lines denote 75% positive expression as a general threshold for MN assays. **E.** Dot plot of gene expression over time in culture (d0-d28). hNIL-iMNs show a marked decrease in expression of markers of pluripotency (NANOG, SOX2, POUF51), and an increase in key markers of iMN identity and maturity (MNX1 (HB9), ISL1, SYP, SNAP25). hNIL-iMNS rapidly exit the progenitor phase, as indicated by the decrease in expression of neural progenitor markers (SOX1, PAX6, OLIG2). **F,G**. Characterization of spontaneous activity (G) and weighted mean firing rate (H). The weighted mean firing rate is defined as the total number of spikes divided by the length of recording of electrodes with activity greater than the minimum spike rate (“active electrodes”). **H**. Activity measured in response to treatment with kainic acid in co-cultured neurons with astrocytes. Error bars represent standard error of the mean. A two sample t-test was used to compare between groups, with a significance level of p = 0.05.

### ML on high-content imaging for discovery of novel disease phenotypes for screening

iPSC-derived motor neurons have been reported to mislocalize TDP-43 under ectopic chemical perturbation (Markmiller et al., 2021; Thams et al., 2019), and in both familial and sporadic ALS patients (Fujimori et al., 2018; Klim et al., 2019); however, phenotypic screens on this low-dimensional signature (mislocalization) have not yielded successful ALS therapeutics (Streit et al., 2022). Our platform uses rich, high-content imaging (HCI) combined with hypothesis-independent ML approaches to uncover novel, ALS-relevant, *in vitro* phenotypes. We measure our diverse cell lines using HCI with channels that span both core morphology and the disease-specific TDP-43. High content imaging provides vast amounts of information on cell biology, most of which is lost upon reduction to one-dimensional phenotypic assays such as TDP-43 mislocalization. We leverage the power of these modern ML methods to learn a foundational representation — an embedding – that reflects the key morphological changes that manifest across these diverse, disease-relevant cell lines.

We show that the combination of HCI and ML is surprisingly powerful, creating an embedding that captures a range of disease-relevant biological phenotypes. In particular, we show that an embedding learned solely using core cell morphology channels accurately predicts TDP-43 mislocalization. Moreover, by measuring bulk transcriptomic data aligned to our imaging datasets, we are able to impute RNA expression from these image embeddings, capturing with high accuracy ALS-relevant transcriptional differences between mutant and WT cell lines. Finally, we show that the same embedding can be used to define a “disease axis” – an ML-derived quantitative score that separates healthy (WT) from diseased (mutant) cells. Our disease axis translates across diverse donor backgrounds, and is much better powered than the TDP-43 phenotype on its own. This provides the foundation both for interrogating mechanisms of disease pathogenesis within and across fALS genetics, and for an *in vitro* screening campaign to discover novel, disease-modifying interventions.

## Results

### Isogenic lines created and differentiated to highly efficient cells of lower motor neuron identity

To reduce non-ALS-related contribution of genetic background on phenotypic signals in phenotypic discovery, we established isogenic lines for donor-specific comparison. For each of the eight fALS gene mutants, we engineered four donors to knock-in (KI) the fALS causal variant or, in the case of TBK1, to create a full knockout (KO). Some lines were removed during editing due to viability issues. Upon edit confirmation, some lines were noted to reflect homozygous mutations knocked-in that were not representative of the majority of patients with the variant (**Table 1**). Introduction of intronic C9ORF72 HRE into these WT) lines was challenging due to size and stability limitations. To address this issue, C9ORF72 patient lines were instead corrected (C9ORF72_corr) through non-homologous end joining (NHEJ) as previously described (Sareen et al., 2013), and were confirmed with both repeat primed PCR (RP-PCR) and fragment analyzer techniques (**Supplemental Figure 1.A**).

**Table 1:** Patient and Engineered iPSC Lines. Parent iPSC Cell lines were onboarded from CIRM (South San Francisco, CA), Cedars Sinai Medical Center (Los Angeles, CA), and Answer ALS that is also sourced from Cedars Sinai Medical Center. Indicated cell lines originally from CIRM were edited at Synthego. C9 Corrected lines and two ectopically expressed TDP-43 cell lines: overexpressing (OE) TARDBP and TARDBP nuclear location signal mutant (NLSm) were edited at insitro. All lines were used in building of assays and formation of the embedding models as described. Anchor analyses and linear classifiers used subsets of these lines per experiment.

| Insitro Cell ID | Donor ID | Cell Line Edit | Clone | Zygosity | iPSC Line Source |
| --- | --- | --- | --- | --- | --- |
| Cins1510 | Dins390 | C9ORF72 | A1 | het | Cedars Sinai Medical Center |
| Cins2178 | Dins522 | C9ORF72 | A1 | het | Answer ALS |
| Cins2179 | Dins523 | C9ORF72 | A1 | het | Answer ALS |
| Cins2183 | Dins527 | C9ORF72 | A1 | het | Answer ALS |
| Cins2186 | Dins530 | C9ORF72 | A1 | het | Answer ALS |
| Cins2188 | Dins532 | C9ORF72 | A1 | het | Answer ALS |
| Cins2189 | Dins533 | C9ORF72 | A1 | het | Answer ALS |
| Cins912 | Dins603 | C9ORF72 | A1 | het | Answer ALS |
| Cins913 | Dins604 | C9ORF72 | A1 | het | Answer ALS |
| Cins917 | Dins605 | C9ORF72 | A1 | het | Answer ALS |
| Cins1509 | Dins390 | corr_C9ORF72 | C3 | NA | Cedars Sinai Medical Center |
| Cins1577 | Dins603 | corr_C9ORF72 | 10 | NA | insitro |
| Cins1579 | Dins604 | corr_C9ORF72 | 3 | NA | insitro |
| Cins1588 | Dins605 | corr_C9ORF72 | 18-2 | NA | insitro |
| Cins1069 | Dins022 | DCTN1-R1101K | B8 | hom | Synthego |
| Cins999 | Dins023 | DCTN1-R1101K | A11 | het | Synthego |
| Cins1086 | Dins025 | DCTN1-R1101K | A2 | hom | Synthego |
| Cins1355 | Dins032 | DCTN1-R1101K | B8 | hom | Synthego |
| Cins991 | Dins022 | FUS-R521C | G5 | het | Synthego |
| Cins992 | Dins023 | FUS-R521C | E2 | het | Synthego |
| Cins1005 | Dins032 | FUS-R521C | H1 | het | Synthego |
| Cins1354 | Dins022 | SOD1-A5V | H7 | hom | Synthego |
| Cins1001 | Dins023 | SOD1-A5V | E3 | het | Synthego |
| Cins1088 | Dins025 | SOD1-A5V | A8 | het | Synthego |
| Cins1496 | Dins032 | SOD1-A5V | C9 | het | Synthego |
| Cins967 | Dins022 | TARDBP-G295S | D11 | het | Synthego |
| Cins972 | Dins023 | TARDBP-G295S | E3 | het | Synthego |
| Cins973 | Dins023 | TARDBP-G295S | H7 | het | Synthego |
| Cins1091 | Dins032 | TARDBP-G295S | F6 | het | Synthego |
| Cins1090 | Dins032 | TARDBP-G295S | E12 | het | Synthego |
| Cins969 | Dins022 | TARDBP-M337V | F8 | het | Synthego |
| Cins974 | Dins023 | TARDBP-M337V | D8 | het | Synthego |
| Cins975 | Dins023 | TARDBP-M337V | F9 | het | Synthego |
| Cins1066 | Dins032 | TARDBP-M337V | H3 | het | Synthego |
| Cins1805 | Dins022 | VCP-R155C | D7 | het | Synthego |
| Cins1063 | Dins023 | VCP-R155C | D1 | het | Synthego |
| Cins1049 | Dins032 | VCP-R155C | E1 | het | Synthego |
| Cins1359 | Dins022 | TBK1-KO | B5 | hom | Synthego |
| Cins1361 | Dins023 | TBK1-KO | E7 | hom | Synthego |
| Cins1363 | Dins023 | TBK1-KO | E4 | hom | Synthego |
| Cins1540 | Dins032 | TBK1-KO | D5 | hom | Synthego |
| Cins1932 | Dins022 | mock-WT | A3 | NA | Synthego |
| Cins1933 | Dins022 | mock-WT | D2 | NA | Synthego |
| Cins2198 | Dins023 | mock-WT | B2 | NA | Synthego |
| Cins1967 | Dins025 | mock-WT | C2 | NA | Synthego |
| Cins1966 | Dins025 | mock-WT | B5 | NA | Synthego |
| Cins1969 | Dins032 | mock-WT | B1 | NA | Synthego |
| Cins2201 | Dins033 | mock-WT | D3 | NA | Synthego |
| Cins128 | Dins022 | Parent-WT | A1 | NA | CIRM |
| Cins1031 | Dins022 | Parent-WT | A1 | NA | CIRM |
| Cins129 | Dins023 | Parent-WT | A1 | NA | CIRM |
| Cins082 | Dins025 | Parent-WT | A1 | NA | CIRM |
| Cins089 | Dins032 | Parent-WT | A1 | NA | CIRM |
| Cins1034 | Dins032 | Parent-WT | A1 | NA | CIRM |
| Cins084 | Dins033 | Parent-WT | A1 | NA | CIRM |
| Cins818 | Dins267 | Parent-WT | A1 | NA | Answer ALS |
| Cins861 | Dins276 | Parent-WT | A1 | NA | Answer ALS |
| Cins1497 | Dins378 | Parent-WT | A1 | NA | Answer ALS |
| Cins1503 | Dins384 | Parent-WT | A1 | NA | Answer ALS |
| Cins1782 | Dins448 | Parent-WT | A1 | NA | Answer ALS |
| Cins1784 | Dins450 | Parent-WT | A1 | NA | Answer ALS |
| Cins1785 | Dins451 | Parent-WT | A1 | NA | Answer ALS |
| Cins1821 | Dins023 | TARDBP(NLSm) | A1 | NA | insitro |
| Cins1676 | Dins023 | TARDBP(OE) | A1 | NA | insitro |

### Rapid transcription factor-based human motor neuron model at day 10

In a series of exploratory datasets, we defined the use of induced transcription factors (TFs) to specify cell fate for use in phenotypic discovery (Fernandopulle et al., 2018). We used an initial subset of lines (n=3) to develop and optimize the protocol (**Figure 1.B**). To generate the MN-like cells, engineered iPSCs were thawed, maintained, and then induced to express three transcription factors (neurogenin 2 (NGN2), islet 1 (ISL1), LIM homeobox containing 3 (LHX3)) (hNIL) through addition of doxycycline for three days; we then banked these for use as neural progenitor cells (NPC). Vials were then thawed and placed in maturation medium for seven days, for a total of 10 days. Immunostaining confirmed that day 10 (d10) hNIL iMNs consistently expressed MN markers across eight donor lines such as HB9 and TUJ1, and produced ALS-related proteins such as TDP-43 and STMN2 (**Figure 1.C,D**). Next, to characterize the maturation of hNIL-iMN protocol over time, we processed two donor cell lines for bulk RNA-sequencing at nine time points ranging from d0 to d28. (**Figure 1.E**). Motor neuron marker expression was consistent with directed MN differentiation protocols (Hu & Zhang, 2010). Notably, genes critical to lower MN function and identity were expressed rapidly at seven days in culture (d10 from iPSC) (**Figure 1.F**). This included increased expression of MN identity genes (MNX1, SLC18A3, CHAT) and neuronal function genes (SYP, SNAP25), and decreased expression of pluripotency genes (NANOG, SOX2, POU5F1). In order to further elucidate the developmental identity of hNIL-iMNs, we did a comparative analysis with pseudo-bulk data generated by Rayon et al. (Rayon et al., 2021) to estimate the Carnegie stage (CS) of cells over different time points. The Spearman coefficient between hNIL-iMNs and fetal MNs from various developmental stages demonstrated a concordance between the *in vivo* CS of brachial and thoracic origin and the time in culture for hNIL-iMNs. Specifically, we found that hNIL-iMN at d3-4 had the highest correlation with CS14 (max corr=0.65); d8-d18 hNIL-iMN had the highest correlation with CS17 (max corr=0.73); and d15-d28 hNIL-iMN had the highest correlation with CS19 (max corr=0.67) (**Supplemental Figure 1.C**).

We next assessed hNIL-iMNs for neuronal excitability with multi-well multielectrode array (MEA) assays. Viable cells exhibited spontaneous action potentials, with an increasing mean firing rate (Hz) over time in culture (**Figure 1.G,H**). We observed extracellular action potentials as early as d2-d5 with a mean firing rate of 0.98 ± 0.1 Hz; these increased to 4.08 ± 1.6 Hz by d23. Co-cultures with rat astrocytes led to the emergence of network bursting activity characterized by an increase in both burst frequency and network burst percentage from d2 to d23 (burst frequency: 0.01±0 - 0.09±0.02 Hz; network burst percentage: 0-64% ± 15 %; see **Supplemental Figure 1.C-E**). Notably, hNIL-iMNs exposed to the glutamate agonist kainic acid exhibited significant increases in network burst potential and synchrony index, providing evidence for functional relevance to essential inputs from corticospinal upper MNs (**Figure 1.I**). Taken together, these data provide insight into the neuronal function, homogeneity, and differentiation dynamics of hNIL-iMNs, further highlighting their applications to ALS disease modeling.

### Automated seeding and dosing of stressors of ALS related pathways

We established an automated seeding, culture, and dosing paradigm that provided fluorescent imaging data and bulk-RNA sequencing data processed from parallel plates in each experiment **(Figure 2.A).** To reduce potential sources of variance in the phenotypic screen, we used automated randomized seeding with interplate and spatial considerations. Isogenic pairs from the same background resided in the same plate with a technical control that was present on all plates. We also used constrained randomization algorithms and multiple-density seeding to de-risk spatial and density effects affecting imaging phenotypes. As a standard input to the high content imaging datasets, we selected a standard staining panel that included both general morphological representation of hNILs (Tuj1, DAPI) and ALS-specific protein localization and expression (TDP-43, STMN2) to enable both classical phenotypic analysis and ML phenotyping. Testing on two WT lines (Dins022, Dins023) displayed expected expression at d10 in culture (**Figure 2.B**).

**Figure 2.**
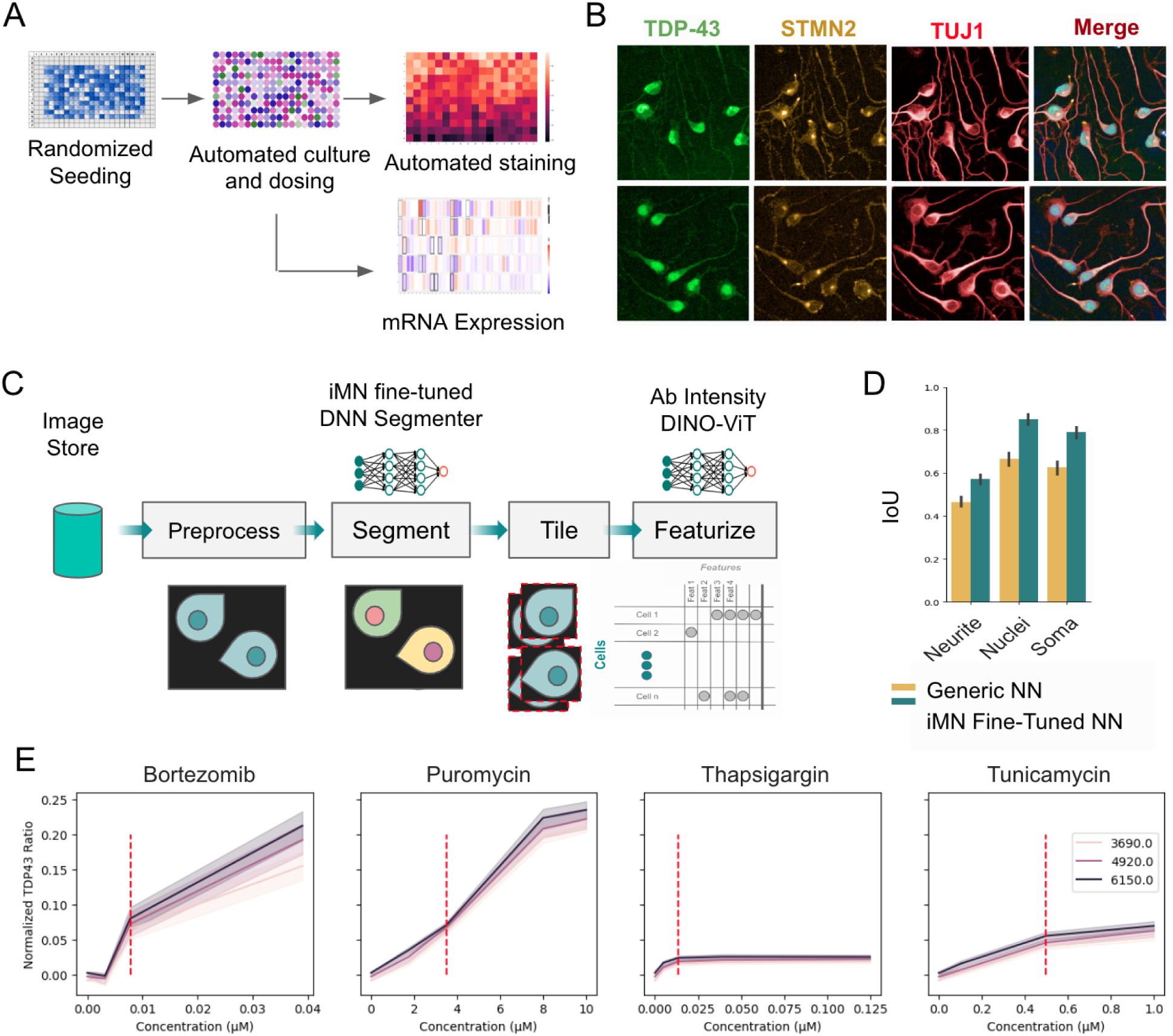
Phenotypic cell platform designed for ML discovery. **A.** Workflow schema of standardized culture, dosing, imaging, and bulk RNA-sequencing of hNILs at d10 for phenotypic discovery. Automated randomized hNIL seeded into 384-well plates were cultured for seven days (d10 total from iPSC). We used digital phase contrast live imaging to QC the cell process at 24 hours post seeding (d4), midculture (d7), and before dosing (d9). At 24 hours before endpoint, we dosed cells with stressors using automated and randomized dosing with an Echo 650 (Labcyte Inc., San Jose, CA). Automated high-throughput staining and imaging generated fluorescence data as input for ML models. Parallel plates from the same randomized cultures were also processed using automated RNA extraction and cDNA normalization for bulk RNA-sequencing. **B.** Representative fluorescent micrograph images of two donor WT hNIL neurons fixed and stained at d10 with a standard panel of antibodies for phenotypic discovery. Nuclei are stained with DAPI (blue in merge). Axonal morphology is represented with TUJ1 (TUBB3) staining. ALS-relevant proteins are represented by TDP-43 localized to the nucleus and cytoplasm and its downstream target STMN2 is found in golgi and neurites. **C.** Schematic of our dataset-generation infrastructure depicting the stages to create a cell-level dataset from raw images. Data stored in the image store are preprocessed, segmented, tiled, and featurized using redun (Insitro, 2021). **D.** An FPN fine-tuned on hNIL-iMN consistently outperforms an FPN trained on a generic dataset, leading to ∼15% higher intersection over union (IoU) and more accurate nuclear, somatic, and neurite segmentation masks. **E.** Dose response curves of TDP-43 C/N in four stressor conditions. Color lines indicate total cells seeded per well. Data are from all WT lines. Red dashed lines indicate selected concentrations used for characterization of mutant lines in heatmaps in Figure 3.B and Supplementary Figure 3.B,E. The shaded area represents bootstrapped 95% confidence intervals.

### Standardized neural network mediated segmentation, tiling, and featurization of datasets in discovery campaigns

To segment the hNIL fluorescent imaging data, we used a composable pipeline that allowed us to compute cell-level statistics in a reproducible and scalable manner using redun (Insitro, 2021). The pipeline is organized into the following modules: preprocessing, segmentation, tiling, and feature extraction (**Figure 2.C**). First, raw images from different microscopes are stored in the cloud using a standardized representation. Next, raw images undergo artifact correction. Quality control (QC) metrics are also calculated as previously described (Sivanandan et al., 2023). Feature pyramid networks (FPN, (Lin et al., 2016)) are then trained and evaluated with annotations generated by human annotators to obtain segmentation masks for nuclear, somatic, and neurite regions. Individual nuclei-centered neuron tiles are generated from estimates of cell coordinates and assigned unique identifiers.

We tested this purpose-built segmentation pipeline alongside a generic neural network segmenter, and it showed improved accuracy when compared to human-annotated ground-truth masks (**Figure 2.D)**. We extracted two types of features from single cell tiles: marker intensities in cell regions like the nucleus, cytoplasm, and neurites, providing us with a set of human-defined features; and a machine-learned representation – also known as an embedding – derived by a self-supervised ML model. Our ML pipeline is based on DINO-ViT (Caron, Touvron, Misra, Jegou, et al., 2021), but with considerable optimizations and enhancements tailored to our particular use case; the resulting ALS-tailored iDINOv3 outperforms pre-trained DINO-ViT in terms of disease classification accuracy (see below). We trained iDINOv3 on different sets of channels. By using only DAPI and TUJ1 channels, we were able to obtain features corresponding solely to core cell morphology. By using all available channels (DAPI, TDP-43, STMN2, TUJ1), we generated embeddings that also encompass known disease-relevant biomarkers.

To potentially exacerbate latent genetic weaknesses in the fALS mutant lines, we administered a set of four stressors known to induce ALS-related phenotypes to hNIL cultures at 5pt and 3pt concentration ranges for imaging and bulk-Seq, respectively. Bortezomib inhibits proteasomal degradation, a key pathway relevant to intracellular aggregation of insoluble TDP-43 (van Eersel et al., 2011). Puromycin has been reported to induce TDP-43 translocation and affects RNA translation, which affects TDP-43 function (Markmiller et al., 2021). Tunicamycin potently inhibits the N-linked glycosylation of glycoproteins, while thapsigargin disrupts Ca2+ homeostasis in the endoplasmic reticulum (ER) (Wu et al., 2018), key pathways affected in ALS. We first tested the ability of our model to capture ALS-relevant biological responses by characterizing the effect of chemical stressors on on a subset of five million cells; we measured the TDP-43 cytoplasm-to-nucleus ratio (C/N) and STMN2 intensity in soma and neurites in all WT lines across three different seeding densities **(Figure 2.E)**. We found that TDP-43 C/N was increased in a dose-dependent manner across all stressor conditions. STMN2 within neurites was decreased by all stressors, whereas bortezomib caused an increase in STMN2 in the soma (**Supplemental Figure 2.B, C**). These results indicate that the automated randomized culture system could readily quantify ALS-relevant biology using the segmentation and analysis workflow at the single tile level, and that stressor conditions reproduced ALS phenotypes for TDP-43 translocation and STMN2 reduction as previously described (M. Y. Fang et al., 2019).

### ALS-related phenotypes quantified across fALS mutants with classic phenotypic analysis

To frame our biological model in previously-described ALS pathology, we sought to determine if any of more than 500 unique culture conditions gave rise to mutant line-specific ALS phenotypes in the hNIL cells at d10. Specifically, two previously-described ALS anchor phenotypes were assessed: TDP-43 C/N, and reduction in expression of STMN2, a downstream target of TDP-43 dysfunction (Klim et al., 2019; Melamed et al., 2019). Given potential phenotypic confounds of culture density, we quantified the correlation between endpoint density of WT lines to TDP-43 mislocalization (**Figure 3.A**). When looking both across cell lines and within wells from individual cell lines, TDP-43 C/N ratio was positively correlated with the density of live cells in each well. To account for this correlation in assessing the effect of mutations on TDP-43 translocation, we took two approaches. First, we restricted comparison of TDP-43 C/N between mutant lines and donor-matched controls to wells with similar numbers of live cells using the overlap of 10-90th percentile values of density (**Supplemental Figure 3.A**). To preserve as many included wells as possible, a second approach used a linear model for TDP-43 C/N that included donor and density of live cells as covariates.

**Figure 3.**
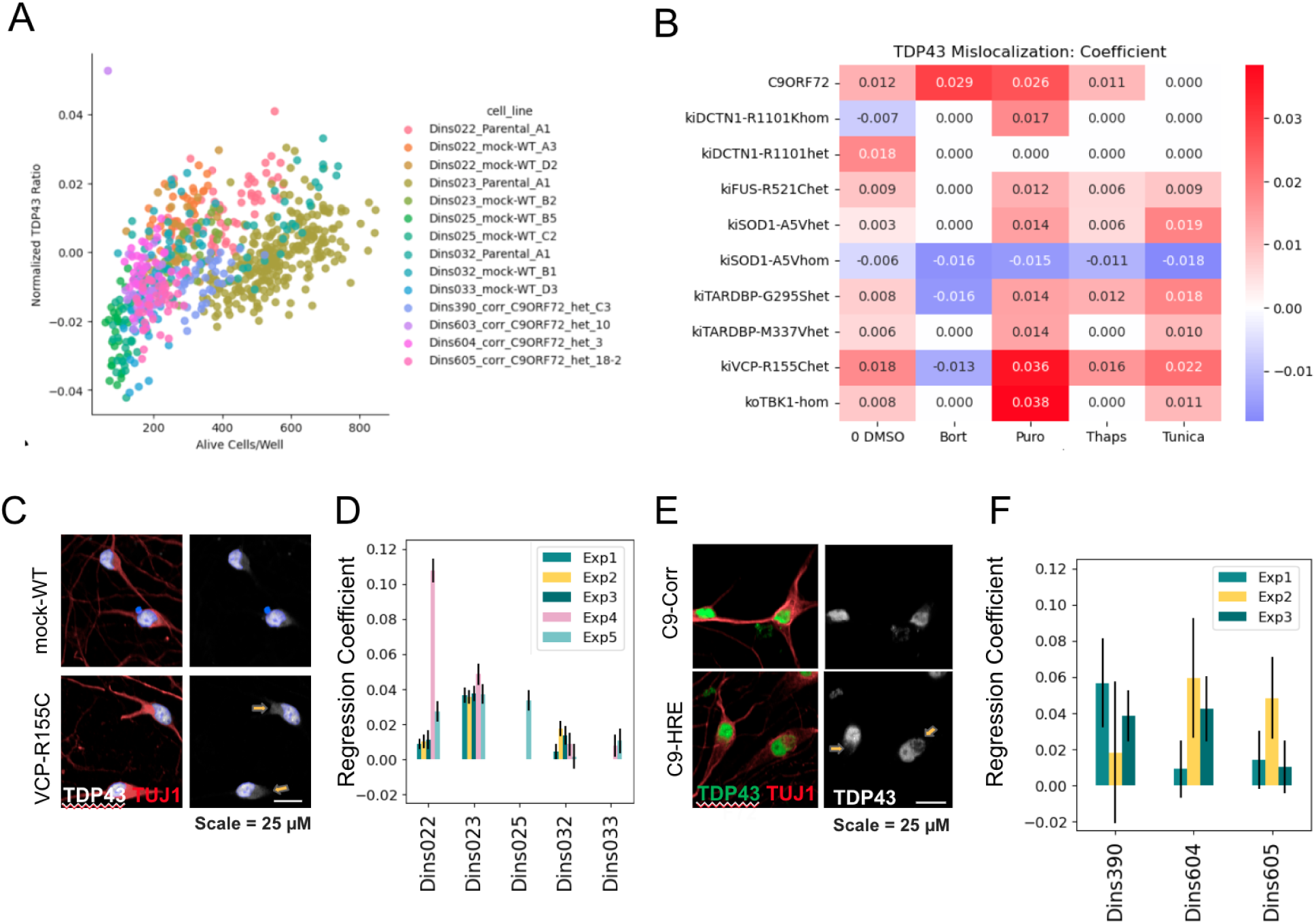
TDP-43 is mislocalized in fALS mutants. **A.** Correlation (R^2^=0.24) between density (Alive in Well) and TDP-43 C/N ratio within and across WT backgrounds indicates that density must be accounted for to determine mutant vs. WT differences in TDP-43 C/N ratio. **B.** Heatmap of TDP-43 C/N mask ratio coefficients using a linear model accounting for density across all wells. Rows indicate tested mutations and columns represent stressor conditions. Red indicates increases and blue indicates decreases. Coefficients with p>0.05 are white. Data are from the primary screen. **C.** Representative fluorescent image of VCP mutant and WT lines. TUJ1 is colored in red, TDP-43 in white, and DAPI in blue in the left panel. TDP-43 is in white and DAPI is in blue in the right panel. Scale bar = 50 uM. **D.** Value and 95% confidence interval of coefficients comparing WT/VCP-R115Chet for across donor backgrounds (x axis values) and experimental repetitions (color). **E.** Representative fluorescent images of C9ORF72 and corrected lines treated with bortezomib. TUJ1 (red) and TDP-43 (green) are in the left panel; TDP-43 staining channel is shown in the right panel. **F.** Value and 95% confidence interval of coefficients comparing WT/VCP-R115Chet for across donor backgrounds (x axis values) and experimental repetitions (color). **G.** C9ORF72 patient lines treated with bortezomib, and coefficients comparing C9ORF72/Corrected TDP-43 translocation in bortezomib.

Using these two approaches, we assessed TDP-43 translocation across seven genes in five stressor conditions in a primary screen. The two approaches yielded similar results for significant differences between WT and mutant lines for primary screen data (**Figure 3.B, Supplemental Figure 3.B**). We found significant (p<0.05) changes in TDP-43 C/N from both models in 13 of 35 conditions. Across a total of 181 comparisons, 121 had significantly altered (p<0.05) TDP-43 C/N ratios between WT and mutant conditions. However, most donor backgrounds did not have consistency in the TDP-43 C/N response (example **Figure 3.B**). Only the VCP-R155C mutation had statistically significant cytoplasmic translocation under the basal (DMSO) condition across 10/11 donor backgrounds and replicate pairs (**Figure 3.C-D)**. Similarly, VCP-R155C had increased translocation in at least 10/11 donor replicate pairs under puromycin, tunicamycin, and thapsigargin stressors, but not under bortezomib. The C9ORF mutation demonstrated a consistent (6/9 comparisons) increase in TDP-43 C/N under bortezomib (**Figure 3.E-F**) and puromycin stressor conditions. No other mutation and stressor condition had >60% consistency across donor and experiment comparisons.

We also observed an STMN2 reduction in fALS mutations across stressor conditions. STMN2 expression intensity within both the soma and neurites was weakly, positively correlated with live cell density (**Supplemental figure 3.D**). Analysis of STMN2 expression intensity within soma and neurites was therefore corrected, as we did for TDP-43 C/N, by using density matching and covariate inclusion (**Supplemental figure 3.E**). Whereas stressor conditions caused a decrease in STMN2 fluorescence intensity overall, we did not observe consistent decreases in STMN2 expression intensity in either soma or neurite compartments when comparing fALS mutant lines to isogenic controls across experimental replicates and stressor conditions (**Supplemental figure 3.F**). This indicates that an STMN2 mutant-specific decrease in expression could not be detected from staining data. Taken together, these results show that, while mutant-specific TDP-43 mislocalization direction was broadly consistent with reports in literature and human patients, the effect size was subtle and sensitive to donor effects.

### Embedding generated from imaging datasets predict molecular features related to ALS

As described above, we generated embedding models trained using distinct image channels, which enabled us to investigate the information content in different biological readouts: nucleus-only (DAPI); nucleus, cytoplasm, and neurite features (DAPI and TUJ1); or all channels, including ALS-relevant biomarkers (DAPI, TUJ1, TDP-43, and STMN2).

We tested the effectiveness of embeddings in capturing ALS-relevant biological information in a series of supervised learning tasks. For this, we used a subset of fALS mutants that had at least three isogenic donor pairs: C9ORF72; SOD1(A5V); TDP-43(G295S); TDP-43(M337V); and VCP(R155C). The first test predicted in-well TDP-43 C/N based upon embeddings from only DAPI and TUJ1 image channels, and hence did not contain TDP-43 information. For each mutant line, we randomly chose 100 wells and divided the single cells in each well into 67% training data and 33% testing data. We then trained a ridge regression model (Hoerl & Kennard, 2000) on each dataset, utilizing grid search for hyperparameter tuning of L2 terms. The mean and standard deviations of the Spearman correlations were 0.569 and 0.09, respectively. The model performed best on TARDBP(G295S) (donor Dins023) with a Spearman correlation of 0.65, and worst on SOD1(A5V) (donor Dins022) with a correlation of 0.52. (**Figure 4.A)**, depicts an example of predicted TDP-43 C/N against the actual values for TARDBP(G295S) (donor Dins023). In all cases, the obtained correlations indicate that our imaging embeddings contain TDP-43-related biology, even without its direct measurement. (**Supplemental Figure 4.A)** shows the results for a single classifier using data from all cell lines and mutations, comprising approximately five million cells. The model was trained on a sub-sample consisting of 80% of the samples that were randomly selected 80%, and was tested on the remaining 20%.

**Figure 4.**
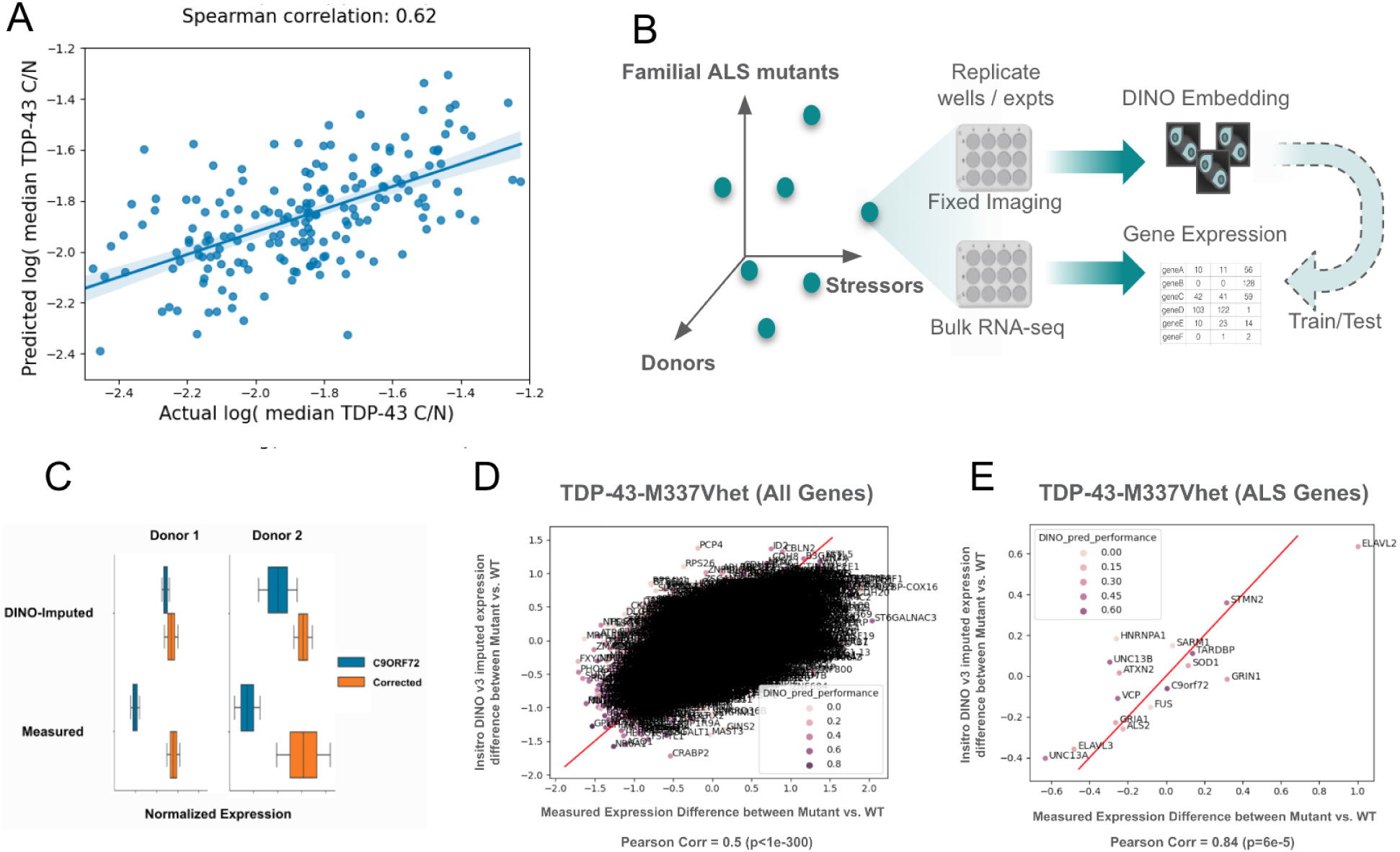
Credentialing tasks confirm imaging embedding contains ALS-relevant information. **A.** Correlation between predicted and actual TDP-43 C/N values derived from morphology embeddings (DAPI and TUJ1). Each point is an individual TARDBP(G295S) cell. Spearman correlation 0.62. **B.** RNA-seq imputation from morphological embeddings. For each condition and experimental batch, we collected fixed-image and bulk RNA-seq data from separate wells. To assess the biological relevance of the morphological embeddings, we examined if they were predictive of RNA-seq from the same condition despite mismatch at cell, well, or plate levels. **C.** Morphological embedding reproduces STMN2 expression decrease in C9ORF72 donors. The STMN2 expression z-score of two pairs (left and right) is shown in the C9ORF72 mutant line (blue) and the mutation-corrected isogenic line (orange) treated with bortezomib. **D.** Scatterplots that compare the measured (x-axis) and imputed (y-axis) change of expression for ALS genes of all 14,000k genes included in the analysis. Pearson (0.5) correlation between the x- and y-axis are annotated in the figure. **E.** Reduced coverage of plot to show ALS-related genes consisting of known ALS GWAS hits and TDP-43 targets.

Next, we assessed the biological relevance of our ML embeddings by investigating their ability to predict gene expression of cells collected under the same experimental conditions, as measured by bulk RNA-seq (**Figure 4.B**). We selected imaging and RNA-seq data from 40 unique conditions per treatment for sufficient quality for paired analysis. Because the RNA-seq was collected at the level of wells, we had a limited number of measurements available for training the predictive model. For each condition, we calculated the mean morphological embedding of all image tiles and the mean gene expression of all RNA-seq replicates. We removed the components of the RNA-seq data explainable by cell count by fitting a LASSO regression model (Tibshirani, 1996) of gene expression using cell count as the only feature and subtracting predicted from measured expression values. We then reduced the dimensionality of the morphology embeddings by performing principal components analysis (PCA) (Hastie et al., 2009) and extracting the top 50 principal components. Lastly, we trained a multitask Lasso regression model that predicted the expression residual of each gene from the projection of each embedding vector onto the aforementioned principal components. In total, we included 14,000 genes in this analysis after filtering genes with low expression value or low variance. For each treatment, we employed cross-validation to assess the predictive performance. For each cross-validation fold, we held out a donor from training, trained a model on the remaining donors, and evaluated the predictive performance on the held-out donors by the Pearson correlation between the observed and predicted expression evaluated for a given gene. (**Supplemental Figure 4.B)**. Correlation of expression values indicated positive imputation prediction across genes above what could be explained by cell density alone. These results suggested that imputation of imaging data to gene expression was likely driven by underlying biology and not by observed variance in cell density.

To further establish the disease relevance of the gene expression imputation, we examined whether it was able to reproduce the measured transcriptomic changes in fALS mutants. We first examined STMN2 expression in C9ORF72 mutant lines. In both DMSO and bortezomib conditions, we observed that the imputed expression recapitulated an observed decreased STMN2 mRNA expression in two C9ORF72 donors that demonstrated this phenotype (**Figure 4.C)**. This analysis paradigm was then expanded to all familial mutants and genes. For TDP-43-M337V in DMSO, the imputation model achieved a Pearson correlation of 0.5 between the imputed and measured change of expression for all 14,000 genes included in the prediction task (**Figure 4.D**). When further subsetting to an ALS-relevant gene set (**Figure 4.E**), the correlation increased to 0.84. Comparable results were observed for TARDBP-G295, VCP, FUS and SOD1 mutations (Supplementary Figure 5). This indicated that the imputed values from imaging data could represent the measured differences in gene expression caused by the fALS mutation.

### ALS disease axes within and across donor lines

Having gained conviction in the disease-relevant information content of our embeddings, we then created a one-dimensional disease axis that best summarized the transition in phenotypic space resulting from a disease-causing mutation. Such a disease axis is a key enabler in a screening campaign towards disease-modifying interventions. We first trained mutant and WT cells separately within each genetic background. For each pair of donor and mutation, we subsetted the phenotypic dataset to contain both WT and mutant cells, embedded the single cell tiles, and trained a classifier on top of the obtained embeddings to distinguish WT from mutant cells. We compared the three models that utilized different subsets of channels. We trained the classifier on a subset of cells, tested on the held-out cells, and evaluated performance using the “accuracy” metric of Carvalho et al. (Carvalho et al., 2019). As expected, all classifiers had high predictive accuracy when built within a one-donor background. The full four-channel model led to higher accuracy, whereas fewer channels showed a modest loss of accuracy (**Figure 5.A**).

**Figure 5.**
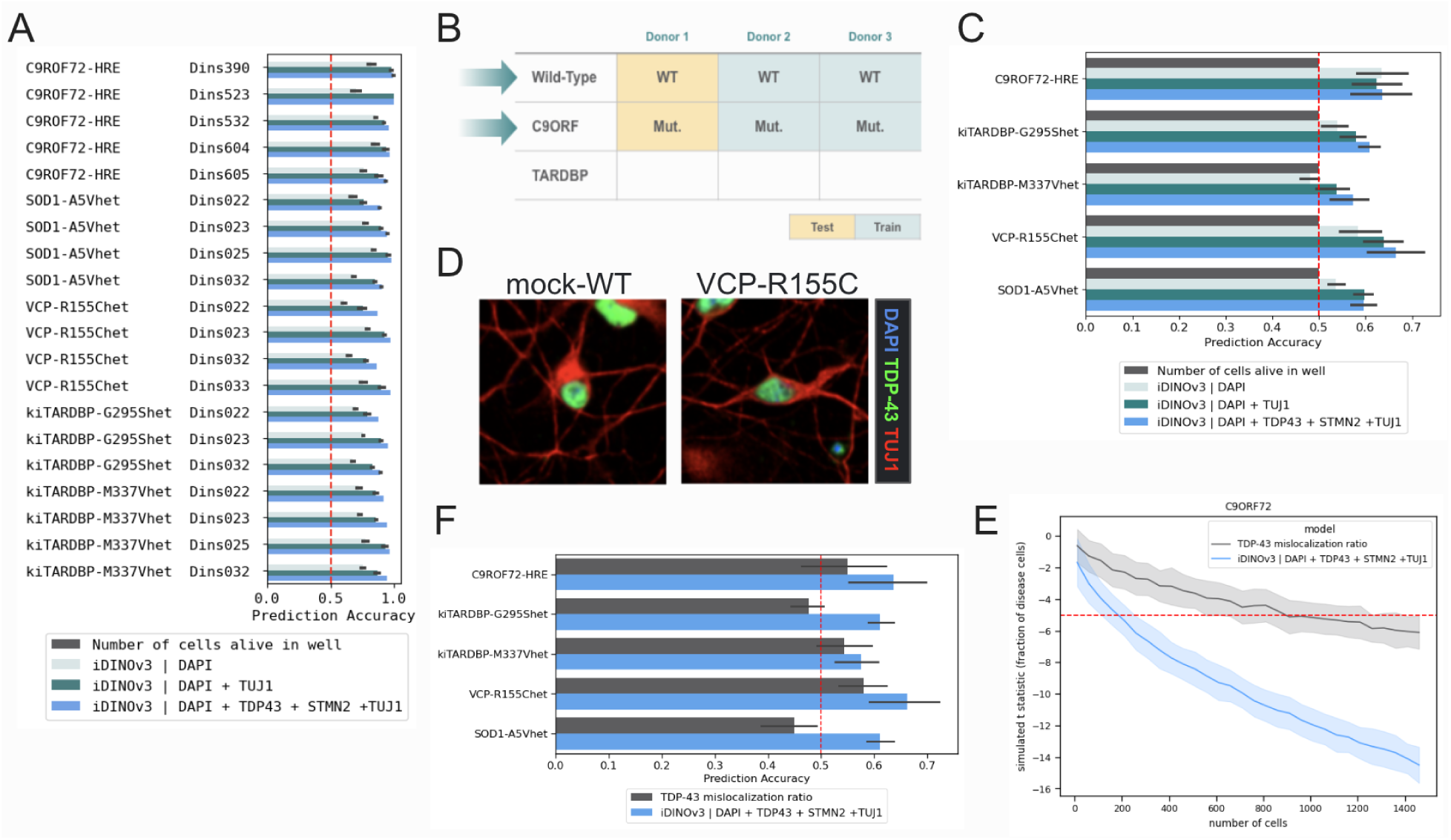
ML methods and results to describe multiple fALS disease classifiers. **A.** Accuracy of linear classifiers differentiating mutant from WT cells within each donor mutant pair. The plot reports the performance of a model trained on well density only (black); DAPI (light green); DAPI and TUJ1 (dark green); and DAPI, TDP-43, STMN2, and TUJ1 (blue). **B.** Schema for the donor hold-out regime. In the example iteration, donor one is held out and the classifier is trained on two remaining donors. For testing, donor one is used to determine classifier accuracy. Then, each donor set is rotated for full test/train representation across three donors within each mutant class. **C.** Mean accuracy of linear classifiers differentiating mutant from WT from donor hold-out regime for each mutant class. The plot reports the performance of a model trained on well density only (black); DAPI (light green); DAPI and TUJ1 (dark green); and all channels DAPI, TDP-43, STMN2, and TUJ1 (blue). **D.** Representative image tiles from extremes of VCP DMSO-treated classifier with all channels, DAPI (blue), TDP-43 (green), and TUJ1 (red). **E.** Cross-donor mutant/WT classifier accuracy on linear classifiers trained using single-feature TDP-43 mislocalization ratio (black), or all channels (blue). **F.** Power analysis of a simulated phenotypic reversion screen for C9ORF72 mutant. The t-statistic is the predicted number of disease cells between groups as a function of the number of cells subsampled, for a simulated dataset composed of 10% mutant and 90% WT cells (group 1) and pure mutant cells (group 2). The red dotted line denotes the significance level, set at five standard deviations. See **Methods** for more details on simulation methodology.

We then moved to the more difficult task of classification across different donors. To reduce the impact of donor-specific genotypic noise, we balanced the number of lines contributing to each mutant and WT class and applied a donor-based normalization method consisting of standardization followed by whitening (see **Methods**). Next, we implemented a donor holdout strategy (**Figure 5.B**): for each mutation, we selected one donor at a time and excluded it from the training data; the remaining donors were then used to train the classifiers. Classifier performance was evaluated using cross-donor validation by testing each model on the donor excluded from its training set and averaging across all permutations. Finally, we calculated the aggregate performance of the classifiers by taking the mean and standard deviations of accuracy of all the models for a given mutation. As per above, we repeated this analysis across different combinations of imaging channels. An initial test performed on C9ORF72 mutants showed a sensitivity of the classifier along differential densities for individual donor pairs. This was confirmed by a classifier trained on density only whose predictions were aligned with cell viability rather than the disease state (**Supplemental Figure 4.C**). To mediate this issue, we followed the same strategy that was used to validate the anchor analysis, that is, stratifying our train and test datasets for wells with similar densities.

The model trained only on well density had an accuracy of approximately 0.5, which demonstrated that the density adjustment procedure did not contribute predictive power as expected. Accuracies for fluorescent channel embeddings ranged from 55% to 70% with DAPI, TDP-43, and TUJ1 channels; kiTARDBP-M337V and VCP had the lowest and highest scores, respectively. Overall, increasing the number of channels improved the classifiers accuracy by ∼10% in distinguishing WT from mutant (**Figure 5.C**). We tested the performance of the classifier compared to using only TDP-43 ratio using the receiver operating characteristic curve (ROC) and found improved sensitivity across all mutants, with some models also enabling increased specificity of mutant status in TDP-43 mutants. We attempted to identify a qualitative biological basis by examining representative cell tiles that were scored with high accuracy (referred to as extreme tiles). For most mutations, this examination failed to reveal appreciable differences visible to the human eye (**Supplemental Figure 6.A-C**). However, VCP mutant extreme tiles contained clear differences in somal and neurite morphology (**Figure 5.D**).

We then tested the extent to which the use of an ML-derived disease axis would provide additional power to detect disease-relevant phenotypic changes, beyond the traditional approach of using an anchor trait such as TDP-43 mislocalization. Specifically, we compared a linear classifier trained on the multi-channel embeddings to a predictor derived solely using the TDP-43 mislocalization ratio (cytoplasmic-to-nuclear mask intensity ratio), and compared the test prediction accuracies. In all cases, the embedding-based disease axes significantly outperformed the anchor trait-based classifiers. In a final analysis of the embedding classifier’s utility in phenotypic prediction and screening potential, we conducted a power analysis using a simulated screen, aimed at detecting a disease-reverting hit (see **Methods**). We compared the anchor phenotype to the all-channel disease axis for C9ORF72 mutants (**Figure 5.F**). To achieve the same power, the embedding-based disease axis required 80% fewer cell tiles than the TDP-43 anchor phenotype.

Taken together, these analyses of imaging embedding classifiers show improvement in the detection of ALS-specifying features across distinct genetic causes of fALS, predict ALS mutant containing lines with morphological information alone, and significantly increase power for therapeutic target screening.

### Classifier predicts morphological neurite deficits in VCP mutants

The visual examination of extreme tiles from the ML disease axis for the VCP mutant lines suggested a neurite deficit phenotype in these mutants. To visually verify this prediction, original sites were reviewed from primary and validation datasets and neurite degeneration was qualitatively observed at a population level within original fluorescent images (**Figure 6.A**). To provide additional evidence for VCP-mediated neurite dysfunction, we assessed paired RNA-sequencing data from DMSO (untreated) VCP wells with gene set enrichment analysis (GSEA). Pathways related to cell cytoskeleton-dependent intracellular transport, dendritic spine morphogenesis, and axon fasciculation were significantly down-regulated in VCP mutant cell lines compared to isogenic control cell lines (false discovery rate (FDR)<0.05; **Figure 6.B,C**). Genes that contributed most to the gene ontology (GO) term ‘regulation of dendritic spine morphogenesis’ enrichment score (ES) were consistently downregulated in VCP mutant cell lines compared to their mock-WT isogenic pairs (**Figure 6.D)**. Thus, in the context of the VCP mutation, the ML model identified a significant biological consequence that was corroborated across both imaging and transcriptional modalities. As of now, other mutant classes require deeper investigation to understand the imaging signature underlying the model’s ability to distinguish mutant from WT lines.

**Figure 6.**
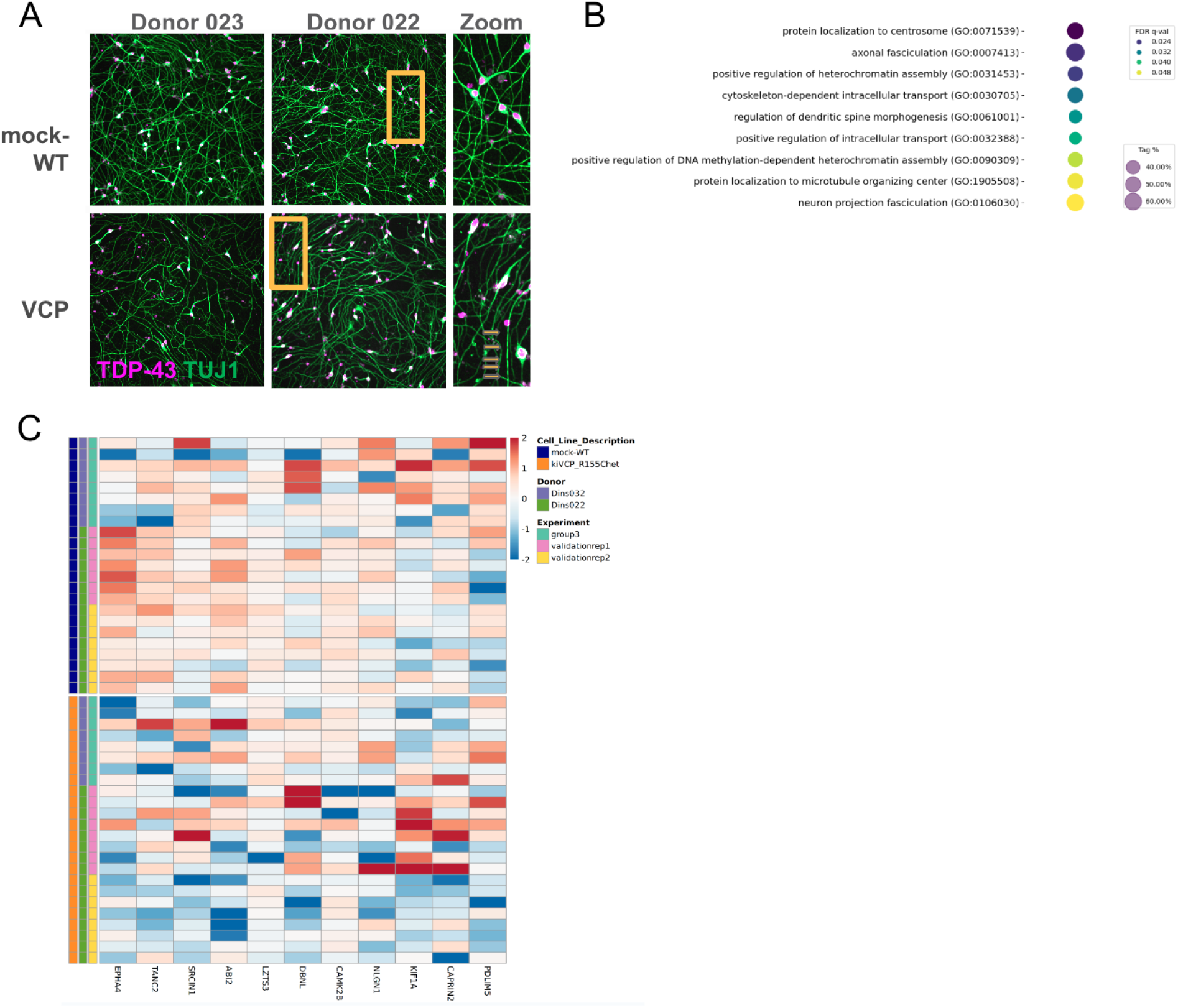
Linear classifier predicts VCP mutant specific neurite dysfunction. **A.** Representative imaging site micrograph of d10 hNIL from two donor pairs (Dins023 and Dins022). Gold boxes denote the spatial area of zoom on the right panel. Yellow arrows denote degradation of neurites observed in VCP mutants. TDP-43 staining is in purple, TUJ1 is in green. **B.** GSEA analysis of genes significantly downregulated in the VCP mutant cell lines are associated to GO terms related to neurite morphology, intracellular transport, and heterochromatin regulation (FDR q-val<0.05; Tag %: percent of GO term genes differentially expressed in our dataset). **C.** Genes that contributed most to the GO term ‘regulation of dendritic spine morphogenesis’ enrichment score (ES were consistently downregulated in VCP mutant cell lines compared to their mock-WT isogenic pairs. The blue-to-red colormap represents log counts per million reads (CPM) expression, clipped to ± 2. n=2 donors and three experiments.

## Discussion

Many diseases, particularly ALS, are difficult to understand and treat because several genes, spanning a broad range of biological processes, contribute to pathogenesis. Traditional models of ALS have often adopted reductionist perspectives which, while valuable, rarely encapsulate the multifaceted nature of such diseases. To counter this limitation, our approach has centered on constructing disease models that allow exploration of this diversity both genetically (through the use of different donor backgrounds) and environmentally (through the use of cell stressors). Development of iPSC-derived patient lines provides an opportunity to study this disease in a more representative, patient-specific manner (Omole & Fakoya, 2018). This approach can be challenging; much of the phenotypic variability observed in different iPSC line backgrounds is likely not directly related to ALS pathogenesis. This complexity is further exacerbated by the fact that the anchor phenotypes such as TDP-43 mislocalization typically used in disease models of ALS patient genetics are fairly subtle (Hawrot et al., 2020; Sances et al., 2016), or not significant (Bilican et al., 2012; Klim et al., 2019). Studies conducted on a large population of lines *in vitro* also did not resolve ALS-specific signatures (Workman et al., 2023). This may be attributed to weak signal that is diminished within the noise. Despite these limitations, previous work has demonstrated that ML classification can predict FALS origin in a small number of VCP mutant lines (Verzat et al., 2022).

To mitigate these challenges and enhance the fidelity of our ALS disease models, we have employed several strategies:

1. Use of isogenic lines, which allows us to isolate the contribution of a specific genetic mutation while controlling for other factors. Moreover, our use of multiple donors allows us to rigorously test the extent to which phenotypes are reproducible across donor backgrounds, thereby controlling for artifacts of specific genetic backgrounds. To our knowledge, our research of multiple mutations across several genetic backgrounds represents the use of the largest set of isogenic lines to date to study any human disease.
2. Development of carefully designed, automated protocols, which have been pivotal in reducing technical heterogeneity, ensuring reproducibility and reliability of results.
3. Implementation of computational methods across our entire workflow, ranging from automated, ML-enabled cell segmentation, to normalization for critical artifacts such as cell density. These tools are key to enabling robust, accurate analyses of the phenotypic data.

The choice of phenotypes measured in disease models is crucial for understanding disease progression. Traditional ‘anchor phenotypes’ in ALS showcase limited cellular changes. However, ALS pathophysiology is highly heterogeneous, with multiple possible trajectories leading to motor neuron death. Anchor phenotypes of ALS are also poorly detected in early-development of model cells; this is not surprising for a disease in which clinical manifestations may not be apparent until decades after disease onset. To address these challenges, we have embraced broad, unbiased imaging measurements and machine learning to identify robust, reproducible phenotypic changes that may be imperceptible using traditional disease models. We provide support for the disease relevance of these phenotypes in a number of ways. Firstly, we were able to impute phenotypes such as TDP-43 mislocalization at the single cell level using only core cell morphology. This is a significant achievement, given the subtlety of this phenotype in ALS. Even more promising, we were able to impute transcriptional profiles, including disease-relevant expression changes, solely from imaging data. Notably, we made these transcriptional imputations across donor backgrounds, demonstrating their robustness. The ability to impute such rich, disease-relevant phenotypes shows that there is considerable information relevant to ALS disease state in our embedding of high-content imaging data.

With this embedding in hand, we were able to establish a disease axis that separates “healthy” (WT) cells from “sick” (mutant) cells. A disease axis encapsulates disease-relevant biological changes in a quantifiable manner, and is a critical tool for identifying ways to reverse and/or slow progression of disease phenotypes. We show that our disease axes are robust both to technical artifacts (plate well effects) and to donor background. Importantly, we demonstrate that this approach provides a considerable increase in power and sensitivity over researching traditional anchor phenotypes, and offers the opportunity to screen for therapeutic interventions using far fewer cells.

Our efforts in this project surfaced challenges that we either addressed or earmarked for future research. Our current methods required dealing with considerable phenotypic variability induced by donor background. Donor background also affected phenotypes both directly and indirectly via cell density in wells, and technical across-well variability contributed to screening complexity. While we were able to considerably ameliorate these challenges by combining experimental and computational methods, we believe that the use of pooled formats containing multiple, genetically-diverse cell lines (“cell villages”; (Mitchell et al., 2020; Neavin et al., 2023)) may appreciably reduce remaining variability and enhance our ability to robustly identify phenotypes; this is a direction of future research. We have only begun to understand the richness of information available in our single-cell images. Axes of variation are hard to discern from images alone, but aligning images with molecular data, such as transcriptomics, may elucidate changes in pathways and might provide more interpretable hypotheses. Our current transcriptomic imputations captured variation across multiple biological processes, some of which are clearly disease-relevant. However, we assessed only bulk transcriptomics at the well level, which did not enable imputation at the single cell level. Using other phenotyping approaches, such as single-cell RNAseq or RNA-FISH, might unlock this potential in future studies.

Taken together, our work demonstrates a unique, ML-enabled platform for studying the *in vitro* phenotypic consequences of complex diseases with genetic bases. It allows us to move beyond reductionist model systems to study such diseases across a breadth of genetic and environmental factors. The combination of high-content phenotyping and machine learning has proven to be surprisingly powerful, and it allows us to interrogate such diseases in an unbiased way. We believe that insights from this research and the future use of our ever-improving platform will unlock novel insights about complex disease processes and lead to the development of therapeutic interventions both in ALS and in other diseases.

## Methods

### Engineering of FALS mutant lines with CRISPR

Donor characteristics and representation are listed in **Supplemental Table 1**. Three non-diseased WT donor control lines (Dins022, Dins023, Dins032) were received under material transfer agreement from California Institute for Regenerative Medicine (CIRM), South San Francisco, CA. Whole genome sequencing data was used with ClinVar (Landrum et al., 2018) to determine that cell lines were not classified as “Pathogenic”, did not have indels of >50 base pairs, and had a ClinVar review status of >2 gold stars. For introduction of mutations, we engineered iPSCs by ribonucleoprotein (RNP) clustered regularly interspaced short palindromic repeats (CRISPR) knock-in methodology (Synthego). Each donor-variant isogenic pair generated two clones that were confirmed for zygosity consistent with fALS patients (i.e., heterozygous (het) or homozygous (hom)) by Synthego’s inference of CRISPR edits (ICE). Due to technical limitations of the performance of the iPSCs through this process, (i.e., poor growth or survival post edit), we excluded some donor-variant isogenic pairs. On rare occasions, we created viable cell lines that had zygosity not representative of patients (i.e., SOD1-homozygous for one donor); we included these lines to determine effects of zygosity and annotated them accordingly. Patient lines with C9ORF72 repeats were acquired from AnswerALS (Cedars Sinai, Los Angeles, CA).

### Engineering of C9ORF72 corrected lines

Native C9ORF72 (G4C2)n donor lines (Dins390, Dins603, Dins604, Dins605) were confirmed for having (G4C2)n expansion through repeat prime PCR (RP-PCR) by gel electrophoresis. Repeat corrected lines (C9ORF72_corr) had repeat (G4C2)n removed through non-homologous end joining (NHEJ) technique previously described in iPSC derived motor neurons *in vitro* (Sareen et al., 2013). Guide RNAs (gRNAs) flanking the repeat region via Cas9-gRNA RNP complex were nucleofected in C9ORF72 patient iPSC lines (Ababneh et al., 2020). We then screened iPSC clones using repeat prime PCR (RP-PCR) by gel electrophoresis. To confirm edits, fragment analysis was also used to confirm (G4C2)n removal.

### hNIL differentiation from iPSCs

Human iPSCs were grown in mTeSR Plus medium (100-0276, StemCell Technologies, Vancouver, Canada) on tissue culture treated plates coated with Vitronectin (A31804, Thermo Fisher) and maintained in 5% CO2 incubators at 37 °C. Cell lines were routinely passaged every 5 days by dissociation using Accutase (A1110501, Thermo Fisher) for 10 minutes at 37°C, and seeded in media supplemented with ROCK inhibitor (Y-27632, S1049, Selleck Chem) for 24 hrs following passaging. To generate stable iPSCs for differentiation towards motor neuron cell fate, cell lines were transfected with a the hNIL piggybac transposon vector with two cassettes (VectorBuilder): a TRE (tetracycline responsive element) promoter system to allow for inducible expression of two transcription factors: ASCL1 (Achaete-scute homolog 1) and DLX2 (Distal-Less Homeobox 2), T2A linked to allow for monocistronic expression, and a CAG promoter driving the constitutive of the neomycin antibiotic resistance gene and the Tet-On3G reverse transcription tetracycline transactivator (rtTA). For transfection, cells were incubated in the presence of liposomes containing the hNIL piggybac vector, the PBase expression vector (pRP[Exp]-mCherry-CAG>hyPBase, VB010000-9365tax, VectorBuilder), using Lipofectamine Stem Reagent (STEM00008, ThermoFisher). Selection of successfully transposed cells was initiated 24 hrs after transfection by the addition of Geneticin™ (500ug/ml, 10131027, ThermoFisher). Donor provenance and karyotypic integrity were validated using KaryoStat (ThermoFisher).

To initiate differentiation of cells towards a motor neuron identity, cells were seeded in plates coated with Vitronectin at a seeding density of 100k/cm, maintained in mTeSR Plus media supplemented with Geneticin (500ug/ml) for 48 hrs, and induced for 72hr with hNIL Induction Media, containing: BrainPhys Neuronal Medium (05790, StemCell Technologies), MEM Nonessential Amino Acids (1X, 11140050, ThermoFisher), GlutaMAX(1X, 35050061, ThermoFisher), N2 Supplement (1X, 17502048, ThermoFisher), B-27 Plus Supplement (1X, A3582801, ThermoFisher), Recombinant Human BDNF Protein (1ug/ml, 248-BDB-MTO, R&D Systems), Neurotrophin-3 (1ug/ml, 267-N3-025, R&D Systems), and Doxycycline Hydrochloride (1ug/ml, 631311, Clontech). Progenitor cells were cryopreserved for future use in micronic vials in CryoStorCS10 (07930, StemCell Technologies). For arrayed imaging experiments, cells were thawed, seeded into 384 well plates coated with poly-l-ornithine (P4957, Sigma-Aldrich) and Laminin-511(RL511-175, Peprotech), and maintained until experimental endpoint in hNIL Maturation media, containing: BrainPhys Neuronal Medium (05790, StemCell Technologies), MEM Nonessential Amino Acids (1X, 11140050, ThermoFisher), GlutaMAX(1X, 35050061, ThermoFisher), B-27 Plus Supplement (1X, A3582801, ThermoFisher), Recombinant Human BDNF Protein (1ug/ml, 248-BDB-MTO, R&D Systems), Neurotrophin-3 (1ug/ml, 267-N3-025, R&D Systems), EdU (5-ethynyl-2’-deoxyuridine, A10044, ThermoFisher) and Doxycycline Hydrochloride (1ug/ml, 631311, Clontech). Plates are fixed in 4%PFA, permeabilized in 0.2% Triton, blocked with Superblock Tween 20 before incubating with TDP-43 (1:100), STMN2(1:100) and b-Tubulin (1:1000) overnight at 4C.

### Multi-well microelectrode array plate preparation

96-well Cytoview MEA plates (Axion Biosystems) were used with the manufacturer’s recommended protocol. We added a5 µL droplet of 1X polyethylenimine (PEI) to the center of each well and incubated it for one hour at 37℃. Post-coating, wells were washed with 2X PBS and 1X distilled water and allowed to dry overnight within a biosafety cabinet. Post-drying, we added a 5 µL droplet of 20 µg/ml of human laminin (RL511-175, Peprotech) to the center of each well and incubated for 2 hours at 37℃. Immediately prior to seeding, laminin was removed but the coated region was not allowed to dry.

### Extracellular recordings and analysis

Extracellular recordings were acquired using the Axion Maestro Pro (Axion Biosystems, Atlanta, GA) in combination with Axion 96-well MEA plates. Each well consists of an 8 channel electrode array and 96 total wells, allowing simultaneous extracellular voltage recordings from up to 768 electrodes at a sampling rate of 12.5 Khz. Raw continuous voltage data was filtered using a 1-pole butterworth bandpass filter (300-5000 Hz). Individual spikes were detected from filtered continuous data via an adaptive threshold set at ± 5.5 standard deviations from each channel’s RMS noise. In all experimental conditions, only “active” electrodes were considered for analysis, defined as electrodes exhibiting activity greater than 5 spikes/min. Single-electrode bursts were defined as at least 5 consecutive spikes with an interspike interval of less than 100 ms. Network bursts were defined as at least 10 consecutive spikes with interspike intervals of less than 100 ms and a minimum of 35% electrodes participating in a network burst. All analysis was performed using Axion’s NeuralMetric Tool Software. Graphing and statistical tests were carried out using GraphPad Prism.

### Constrained randomization and automated seeding, dosing, and staining

We applied constraints to known spatial effects that correlate with patterns produced by automated processing (i.e, instrument aspirate or dispense patterns that may arise from automation machinery). We randomized well-to-condition assignments using a mixed integer linear program solver, then used internally-built software to map the solutions into specific transfer list files and Echo 650 (Labcyte Inc., San Jose, CA) liquid handlers. Shortly following in-lab execution, automation machine log files were then programmatically propagated back to our electronic lab notebook (ELN) to ensure metadata provenance in this ELN (Benchling, San Francisco, CA). Day three hNIL neural progenitor cells (NPC) were thawed and seeded into a 384-well Cell Carrier (Perkin Elmer, Waltham, MA), using Certus Flex (Fritz Gyger AG, Gwatt, Switzerland). Culture wells were fed with a Biotek EL406 (Agilent, Santa Clara, CA) for seven days. Twenty-four hours before endpoint, we dosed cultures using an Echo 650. Stressor concentrations for imaging include 0, 0.00128, 0.0032, 0.008, 0.04uM for Bortezomib; 0, 1.8, 3.7, 8.2, 10.1uM for Puromycin; 0, 0.01, 0.1, 0.5, 1uM for Tunicamycin and 0, 0.005, 0.0135, 0.0405, 0.1215uM for Thapsigargin. Stressor concentrations for transcriptomics include 0.00128, 0.008uM for Bortezomib; 1.8, 8.2uM for Puromycin; 0.008, 0.5uM for Tunicamycin; 0.005, 0.0135uM for Thapsigargin. We fixed and stained cells on Prime automated liquid handling system (HighRes Biosolutions, Beverly, MA) and imaged on the Opera Phenix (Perkin Elmer) using a 20x water objective (Perkin Elmer). We randomized the processing order of plates throughout this experimental workflow to reduce plate order effects.

### Image analysis pipeline

We adopted multiple widely-used segmentation models, including an FPN (Lin et al., 2016), a pre-trained Cellpose (Stringer et al., 2021), and a fine-tuned Cellpose on iMN. The trained models leveraged a manually curated and annotated dataset of motor neurons. We extracted features by transforming images into vectors used to train classifiers. We computed two types of features: interpretable, hand-engineered features (anchors); and ML-learned features.

### Anchor assays, TDP-43 mislocalization, and STMN2 intensity assays

TDP-43 cytoplasmic/nucleus ratio (C/N) was calculated by taking median intensity of staining within the cytoplasmic mask divided by median intensity within the nucleus mask. The linear regression for TDP-43 C/N and STMN2 heatmaps was calculated with the following regression equation:

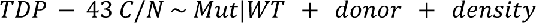

We normalized aggregated values by subtracting the median value of each signal across WT-BTC wells in DMSO to account for plate-to-plate variance. We characterized differences in these signals across mutations by the coefficient of the linear regression either accounting for density in all wells, or with density removed with the dataset that selected for same density comparisons in Supplementary Figure 3.

### Embedding imputation and classifiers

We used self-supervised learning (SSL) training to create embeddings without manual annotations. DINO (Caron, Touvron, Misra, Jégou, et al., 2021), was used to train on our in-house motor neuron datasets referred to as iDINO. The model training took ∼140 hours on eight NVIDIA A100 GPUs. Each cross-validation fold held out a donor from training, and evaluated the predictive performance on the held-out donor by the Pearson correlation between the observed and predicted expression evaluated for a given gene. Our normalization method consisted of standardization followed by whitening (Michael Ando et al., 2017). In the standardization step, we first subtracted the mean (over the training set) vector from each sample and then divided each sample by the standard deviation. In the whitening step, we performed eigenvalue decomposition of the covariance matrix of the data matrix. We then normalized the eigenvalues to have unit variance by taking the inverse square root of the eigenvalues. We ended up with a whitening transformation matrix, W, by multiplying eigenvectors and normalized eigenvalues; we used W to transform our data.

Algebraically, this can be defined as follows. Let *X* denote the original data, *V* be the matrix of eigenvectors, and 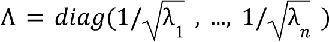 be the diagonal matrix of normalized eigenvalues. The whitening transformation matrix is defined as *w* = *S* * Λ. Thus, the data are transformed as *X*’ = *X* * *w*.

### Power analysis for screening simulation

We simulated a screening target with a 90% reversion rate (i.e., composed of a mixture of 10% mutant and 90% WT cells). Subsampling the results of the linear classifier to different numbers of cells, we then calculated the t-statistic in the resulting fraction of predicted disease cells in the simulated target vs. a pure 100% mutant population, and plotted this t-statistic as a function of number of cells. To achieve a ™5 SD significance level (i.e., high confidence that the simulated target is distinguishable from pure mutant in the healthy direction), the C9ORF72 linear classifier trained on the all-channel model needs approximately 200 cells, whereas the anchor analysis model requires approximately 1000.

### Bulk-RNA processing, sequencing, and DE and GSEA analyses

Day 10 hNIL were used for bulk RNAseq processing. Cells were lysed and RNA was extracted using MagMAX mirVana Total RNA Isolation kit in an automated protocol (Applied Biosystems, Waltham, MA) on the KingFisher Apex liquid handler (Thermo Scientific, Waltham, MA). We QC’d the purified RNA on the Agilent Tapestation 4200, and then normalized to 2 ng total input on the Starlet liquid handler (Hamilton, Reno, NV). We executed bulk RNAseq library preparation using the “Stranded Total RNA Prep, Ligation with Ribo-Zero Plus” protocol (Illumina, San Diego, CA) on the Bravo liquid handler (Agilent). Final cDNA libraries were QC’d on the Tapestation 4200 (Agilent). Samples that passed library preparation were normalized to 10 nM and pooled using Starlet liquid handler. We sequenced pooled final libraries targeting 20 million reads per sample on the NovaSeq 6000 S4 flowcell (Illumina, San Diego, CA).

We trimmed sequencing reads to 30 base pairs and aligned them to the human reference genome GRCh38.p13 (Schneider et al., 2017) using STAR v2.7.5b (Dobin et al., 2013) and to the transcriptome using RSEM v1.3.3 (Li & Dewey, 2011). Gene expression values are presented as log2-transformed counts-per million reads (logCPM), using a pseudocount of 1e-3. We calculated differential expression and gene set enrichment analysis (**Figure 6**) on the raw counts matrices of mutant cell lines vs. mock-WT isogenic controls using DESeq2 v1.34.0 (Love et al., 2014) and GNUparallel (Tange, 2023) with cell culture plate and experiment as covariates. We identified pathways using GSEApy v1.0.5 (Z. Fang et al., 2023). Adjusted p-values and q-values <0.05 were considered statistically significant.

## Supplemental Figures and Data

**Supplemental Figure 1.**
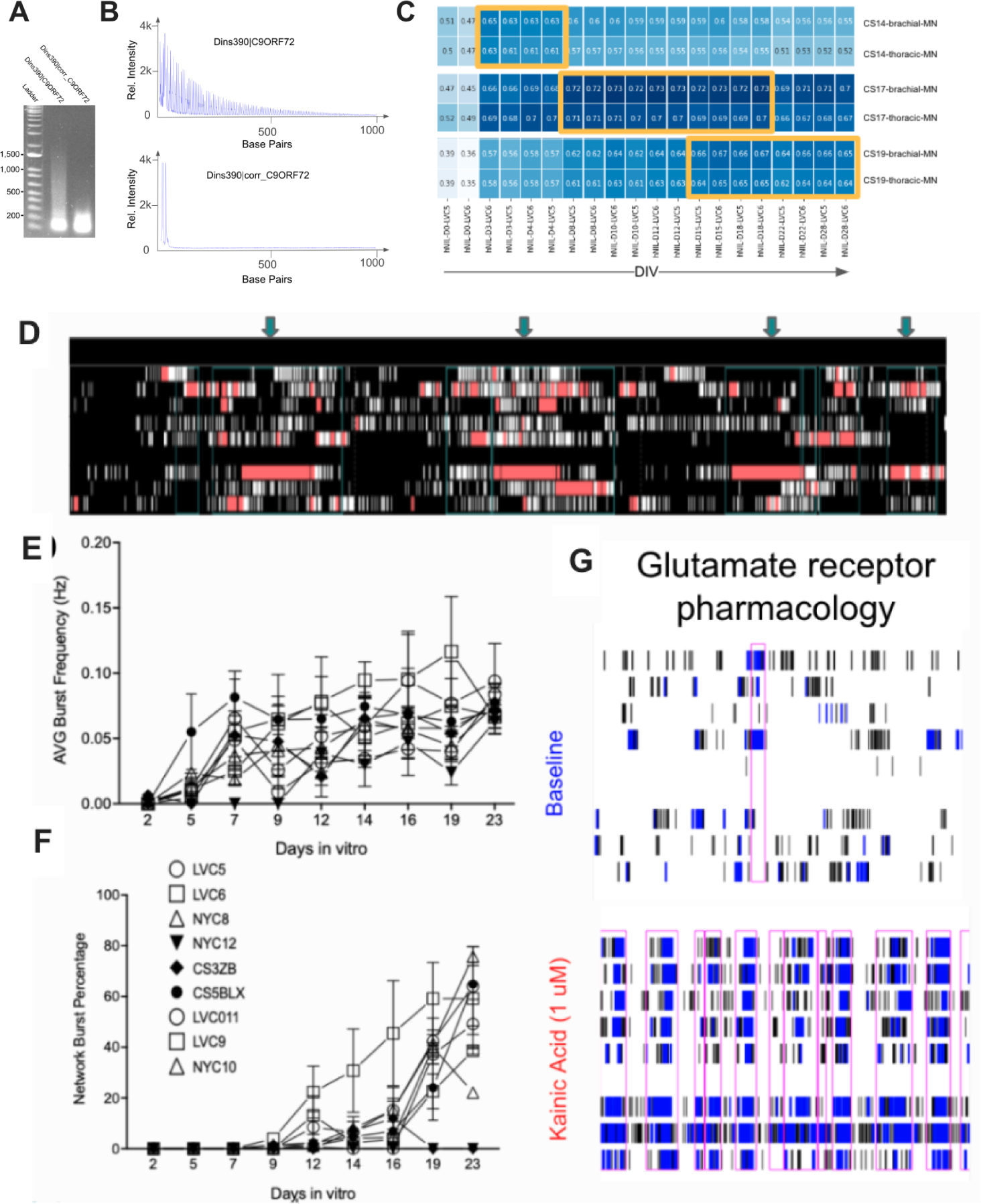
C9ORF72 editing, hNIL developmental Identity, and electrophysiological function with co-culture of astrocytes. **A.** PCR products optimized for C9ORF72 G4C2 repeats are visualized on 1% agarose gels. As an example, for the Dins390 pair, the unedited repeat line has smears at higher molecular weights; the repeat removal line loses the smears. 200 bp, 500 bp, 1000 bp, and 1500 bp DNA markers are labeled in the ladder lane for size estimation. **B.** Repeat number quantification by fragment analysis. We loaded RP-PCR products and standard samples to the fragment analyzer and analyzed them with Geneious software (Geneious, Auckland, New Zealand). Horizontal axis is in base pairs (bp), while the vertical axis is the relative intensity. From the blue electropherogram traces, G4C2 repeat PCR peaks are identified, and converted to repeat length. Repeat numbers (<145) are calculated after calibration with the standard. **C.** Correlation of hNIL-iMN gene expression data with Carnegie stages (CS) of fetal brachial and thoracic MN development. Over the course of time in culture, hNIL iMNs traverse the CS14 to CS19 brachial and thoracic states of fetal development. **D-F.** Characterization of network bursting frequency and network burst percentage after co-culture with rodent astrocytes. **F,G.** Network bursting raster potential in response to kainic acid treatment.

**Supplemental Figure 2.**
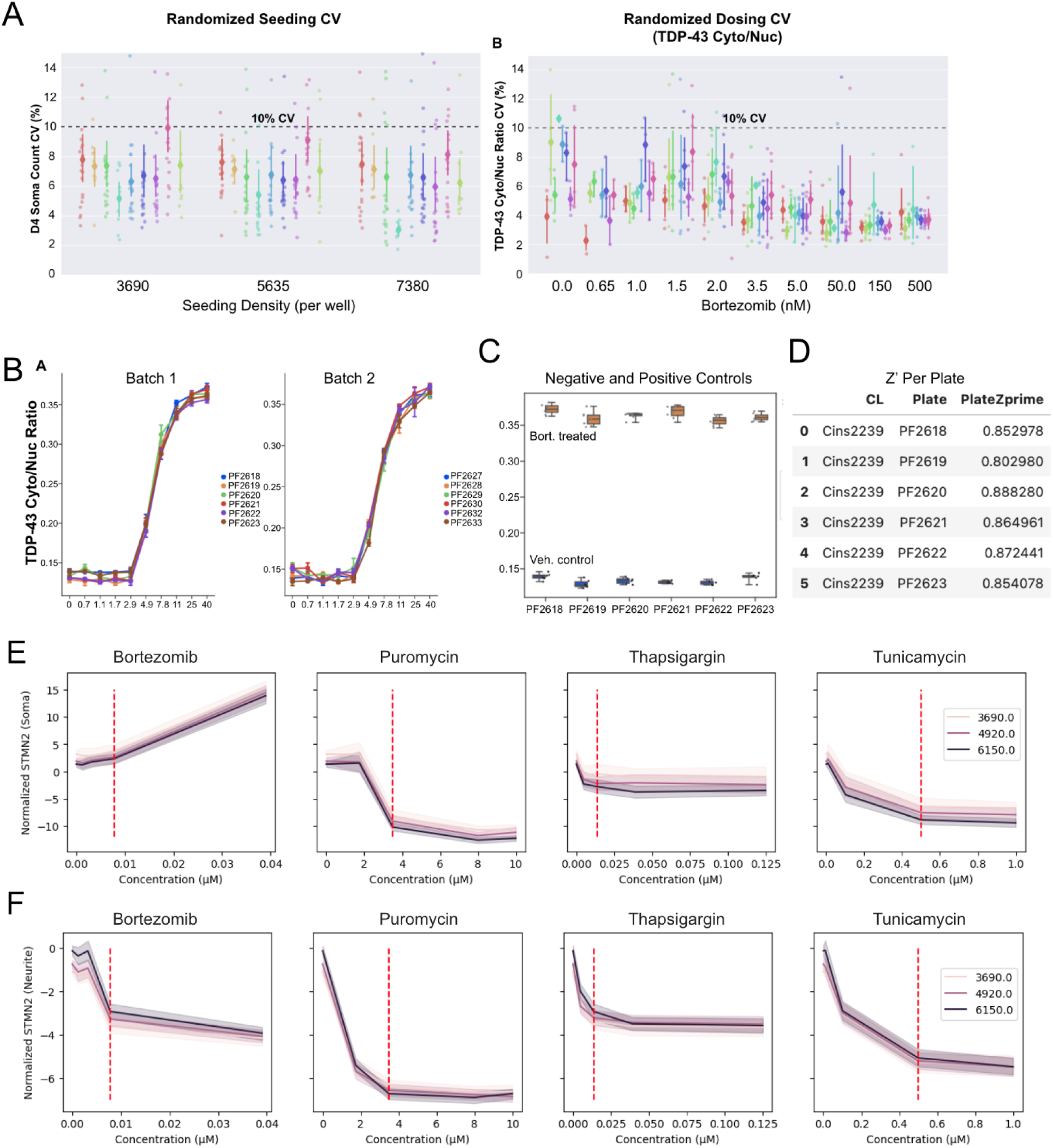
Platform reproducibility and stressor dosing characteristics. **A.** Technical reproducibility of randomized seeding assessed by coefficient of variance (CV) of DPC imaging at d4 (24 hours post plating). Dotted line denotes 10% CV. n=16 plates. **B-C.** Assay reproducibility and screen quality metrics. Technical controls from the fALS pilot screen demonstrated reproducibility of stressor-induced TDP-43 response within and across all plates. **B.** Technical control 1: a 10 point dose response of bortezomib-treated control line in every plate showed high reproducibility. Error bars represent the standard deviation across independent wells. **C.** Technical control 2: six wells each of DMSO and bortezomib-treated (40nM) batch control cell lines randomly distributed in every plate showed low variance and consistent dynamic range within and across plates. **D.** Screening quality metric Z’ calculated using technical control 2 was >0.8 for all plates in the pilot run, meeting success criteria for high reproducibility. **E.** STMN2 intensity within the soma. **F.** STMN2 intensity within neurites. Bortezomib elevates STMN2 expression in the soma, whereas all other conditions decrease STMN2 expression in a dose-dependent manner.

**Supplemental Figure 3.**
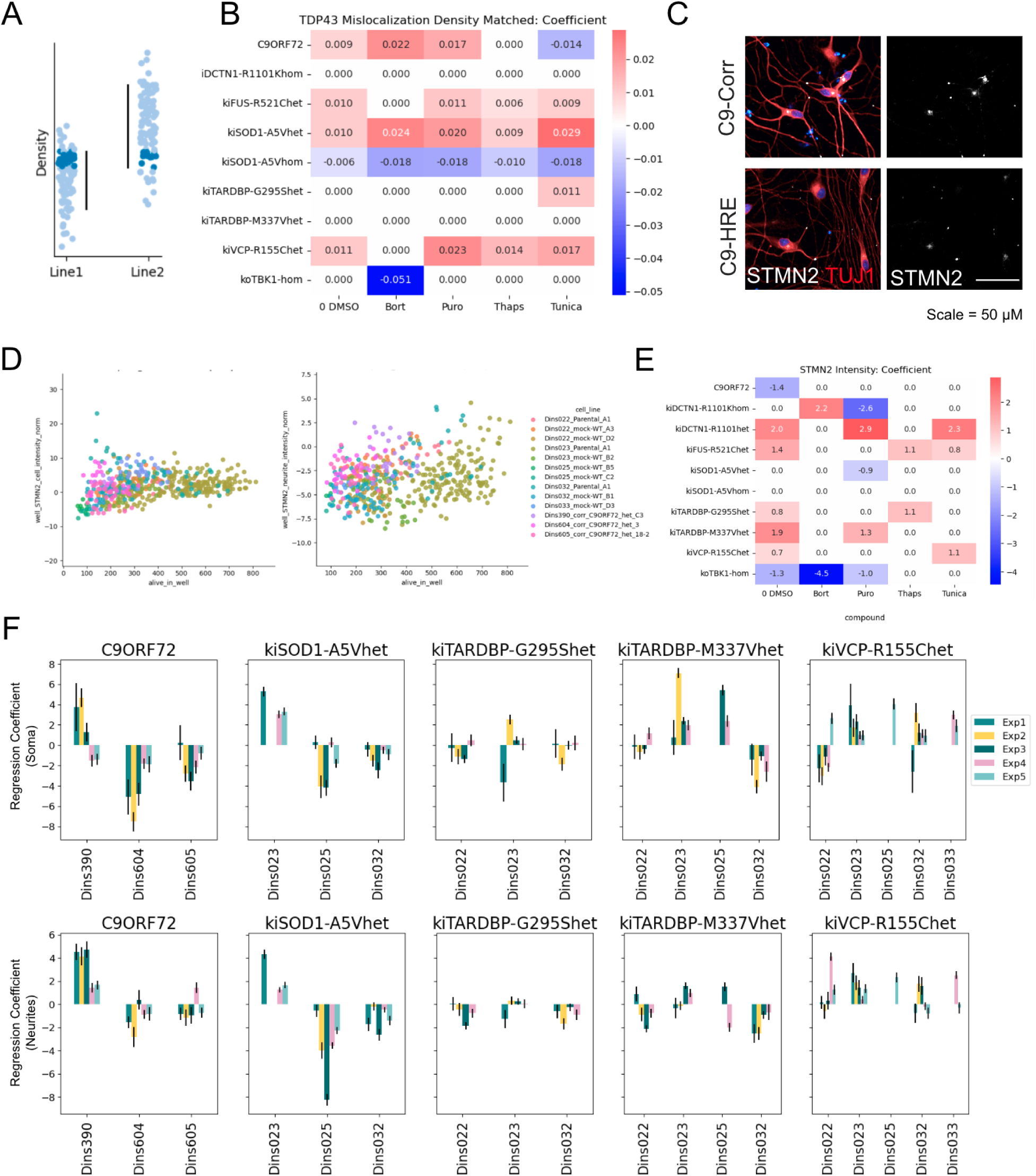
TDP-43 mislocalization density matching and STMN2 staining analysis. **A.** Schematic showing how well replicates from within isogenic WT/mutant pairs were selected, using the overlap of 10-90th percentile values of density. **B.** Heatmap of TDP-43 C/N ratio coefficients from density-matched comparisons (as in B). Rows indicate tested mutations and columns represent stressor conditions. Red indicates increases and blue indicates decreases. Coefficients with p>0.05 are white. Data are from the primary screen. **C.** Representative fluorescent micrograph image of STMN2 staining between C9ORF72 patient (top panel) and corrected line (bottom) in DMSO-treated wells. **D.** STMN2 intensity was weakly correlated in the soma (left, R^2^ = 0.12) and in neurites (right, R^2^ = 0.11) with live cell density. Wild-type cell lines are represented here by color. **E.** Heatmap of STMN2 expression intensity coefficients in the the soma from a linear model accounting for donor and live cell density. Rows indicate tested mutations and columns represent stressor conditions. Red indicates increases and blue indicates decreases. Coefficients with p>0.05 are white. Data are from the primary screen. **F.** Soma (top) and neurite (bottom) STMN2 expression intensity under basal conditions (DMSO) was inconsistently altered by fALS mutations. Each chart shows the value and 95% confidence interval of coefficients from comparing WT/mutant for each mutation (column) across donor backgrounds (x axis values) and experimental repetitions (color).

**Supplemental Figure 4.**
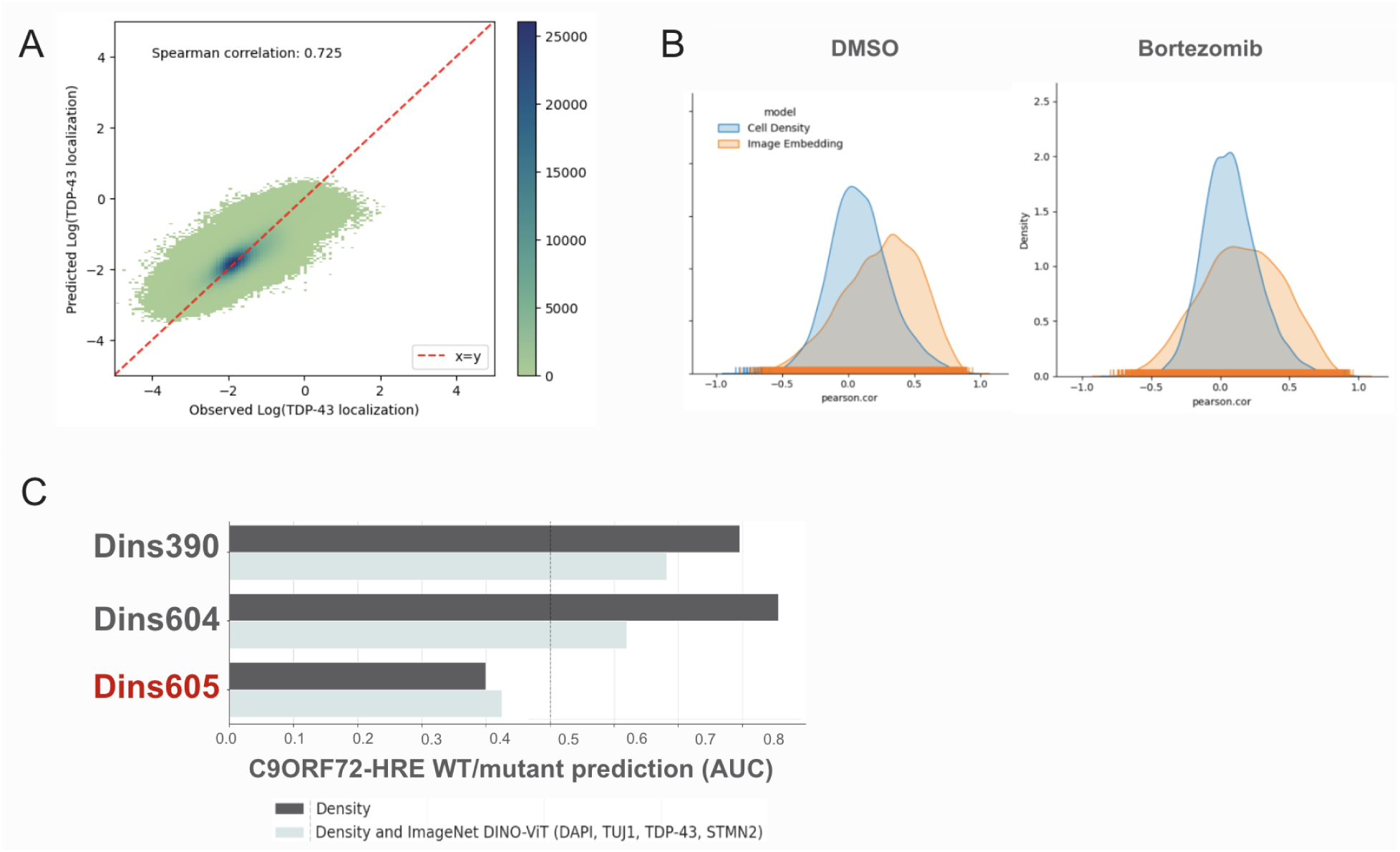
ML classifier density confound and imputation details. **A.** Density histogram of the TDP-43 localization as predicted from DAPI and TUJ1 fluorescence channel embedding, compared to the cytoplasmic/nuclear mask intensity computed from the TDP-43 channel. **B.** Histograms of Pearson correlation between imputed and measured gene expression under DMSO (left) and Bortezomib (right) treatment. The blue histograms are for imputation models using cell density as feature and the orange histograms are for imputation models using image embeddings as feature. **C.** AUC of three different donor pairs of C9ORF72 (Dins390, Dins604, Dins605). Red text denotes a line that had differential density separation that the previous two. Accuracy of linear classifier trained on density showing lower than 0.5 indicates worse than random prediction, indicating density is a strong predictor of overall performance.

**Supplemental Figure 5.**
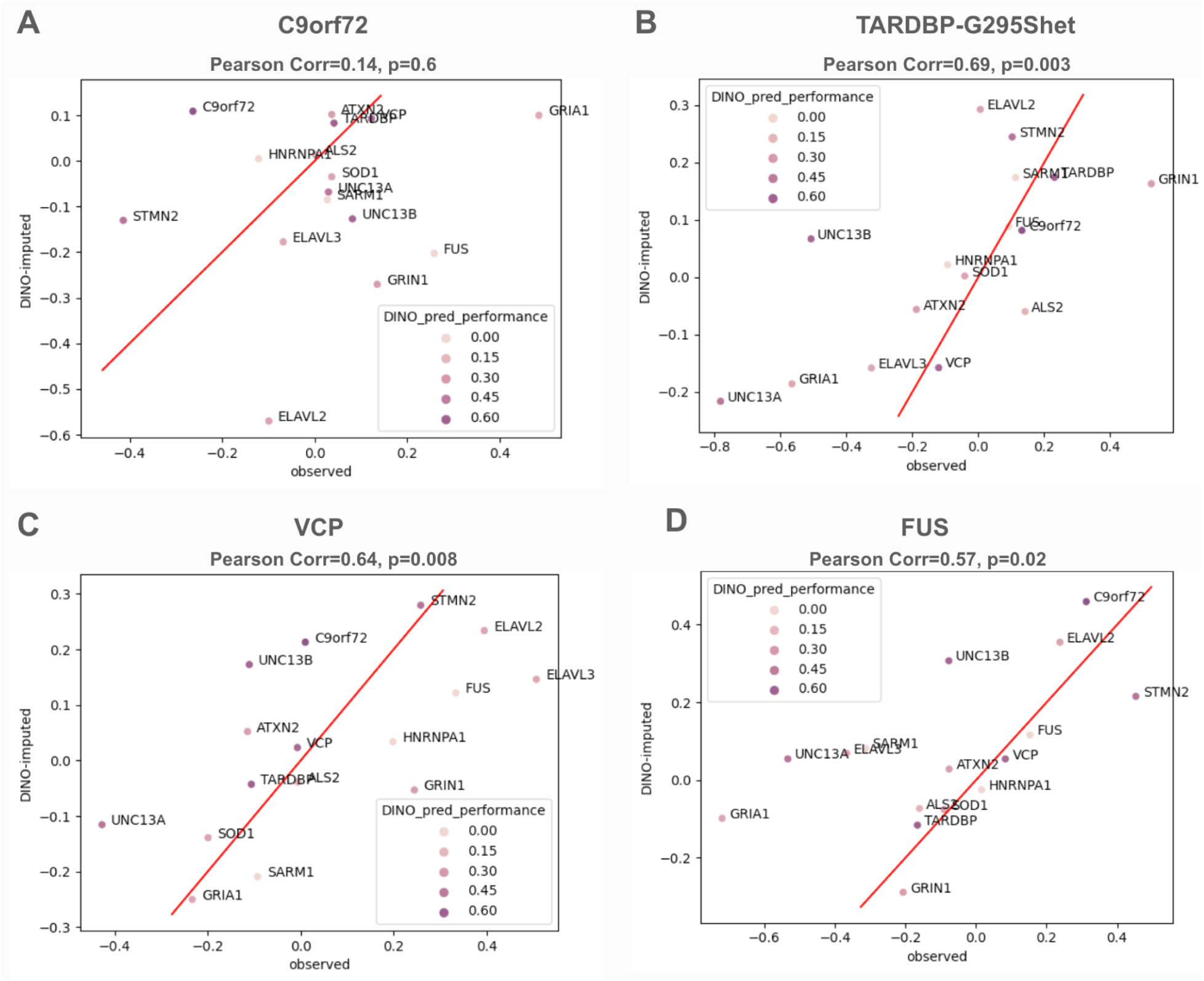
RNA imputation from morphology-derived embeddings for ALS relevant mutations. Genes previously implicated in ALS are shown in the plots. Boxes highlight TDP-43 target genes ELAVL3 and UNC13A. **A.** C9ORF72 **B.** TARDBP-G295Shet **C.** VCP **D.** FUS **E.** SOD1-A5Vhom

**Supplemental Figure 6.**
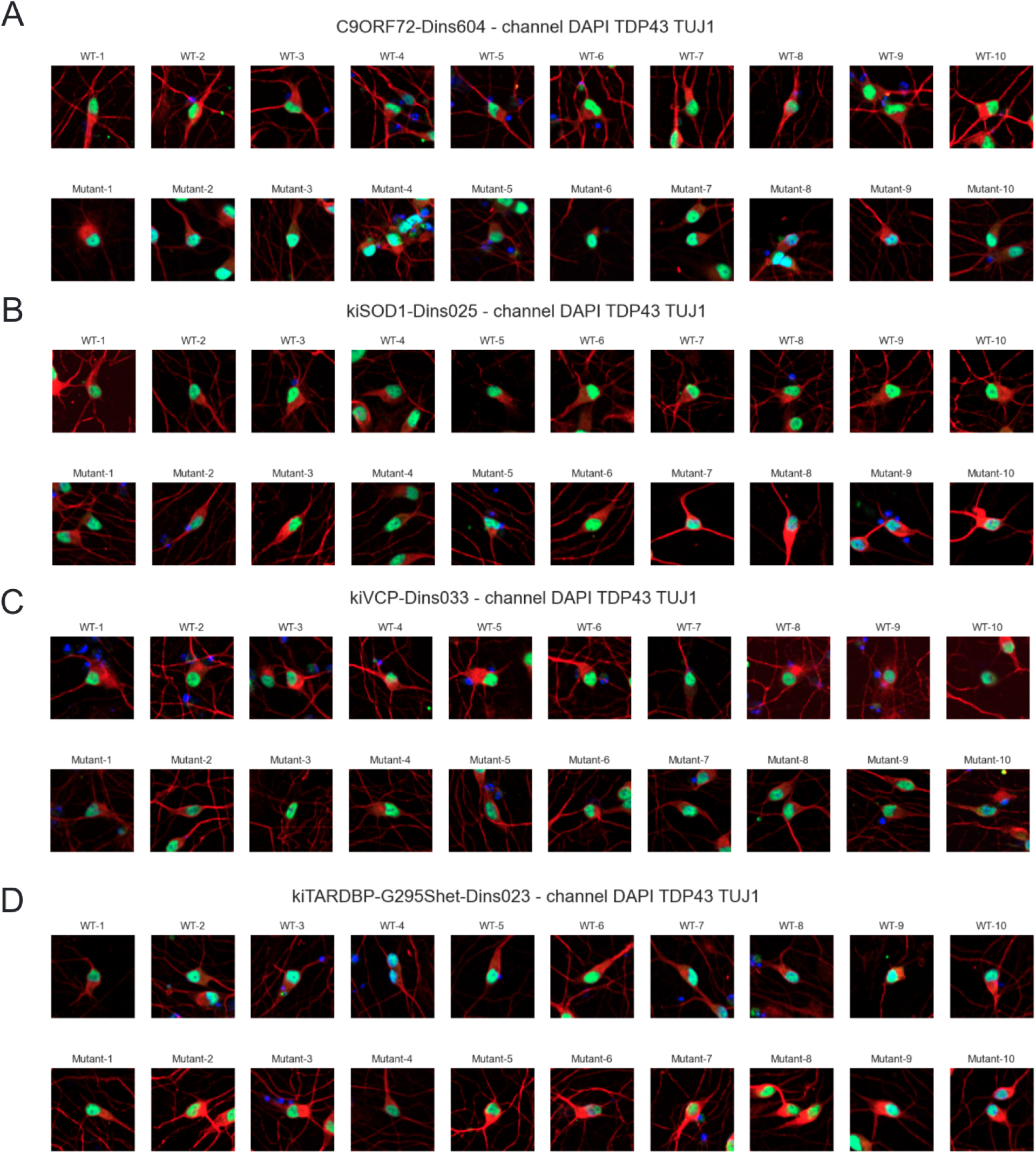
Representative image tiles from extremes of mutant classifiers. **A.** Representative extreme examples of cells from different C9ORF72 classifier. **B.** Representative extreme examples of cells from different SOD1 classifier. **C.** Representative extreme examples of cells from different VCP classifier. **D.** Representative extreme examples of cells from different TARDBP-G295S classifier.

## Acknowledgements

We would like to acknowledge contributors to the insitro platform that was critical in the execution of this work:

Chris Baker, Anthony Ha, Babacar Ndoye, Prateek Chandra, Rob Miller, Kelsey Dexter, Pat Castillo, Madeline Tran, Herve Marie-Nelly, Avantika Lal, Eugeni Vaisberg, Kat Titterton, John Bisognano, Xueya Zhou, Rounak Dey, Eilon Sharon, Audrey Musselman-Brown, Rosey Stone, Max Bates Pooja Prasad, Alicia Lee, Sylvie Gao, Arnab Banik, Karen Lee, Tracy Chan, Blake Simmermon, Liz Ellithorpe, Vincent Luczkow, Ci Chu, Alicia Cuevas, Piotr Kaleta, Anna Shcherbina, Ayla Nelson, Zack Phillips, Brian Young, Meiliang Pan, Heidi Shin, Chungnam Chan, Colm O’Dushlaine, David Conegliano, Sahil Malpotra, Maisha Rashid, Rachel Groth, Sheetal Modi

We would also like to thank Carrie Cizauskas for thoughtful editing contribution and proofing the manuscript.

## References

Ababneh, N. A., Scaber, J., Flynn, R., Douglas, A., Barbagallo, P., Candalija, A., Turner, M. R., Sims, D., Dafinca, R., Cowley, S. A., & Talbot, K. (2020). Correction of amyotrophic lateral sclerosis related phenotypes in induced pluripotent stem cell-derived motor neurons carrying a hexanucleotide expansion mutation in C9orf72 by CRISPR/Cas9 genome editing using homology-directed repair. Human Molecular Genetics, 29(13), 2200–2217.

Akçimen, F., Lopez, E. R., Landers, J. E., Nath, A., Chiò, A., Chia, R., & Traynor, B. J. (2023). Amyotrophic lateral sclerosis: translating genetic discoveries into therapies. Nature Reviews. Genetics, 24(9), 642–658.

Bardy, C., van den Hurk, M., Eames, T., Marchand, C., Hernandez, R. V., Kellogg, M., Gorris, M., Galet, B., Palomares, V., Brown, J., Bang, A. G., Mertens, J., Böhnke, L., Boyer, L., Simon, S., & Gage, F. H. (2015). Neuronal medium that supports basic synaptic functions and activity of human neurons in vitro. Proceedings of the National Academy of Sciences of the United States of America, 112(20), E2725–E2734.

Bilican, B., Serio, A., Barmada, S. J., Nishimura, A. L., Sullivan, G. J., Carrasco, M., Phatnani, H. P., Puddifoot, C. A., Story, D., Fletcher, J., Park, I.-H., Friedman, B. A., Daley, G. Q., Wyllie, D. J. A., Hardingham, G. E., Wilmut, I., Finkbeiner, S., Maniatis, T., Shaw, C. E., & Chandran, S. (2012). Mutant induced pluripotent stem cell lines recapitulate aspects of TDP-43 proteinopathies and reveal cell-specific vulnerability. Proceedings of the National Academy of Sciences of the United States of America, 109(15), 5803–5808.

Brown, R. H., & Al-Chalabi, A. (2017). Amyotrophic Lateral Sclerosis. The New England Journal of Medicine, 377(2), 162–172.

Caron, M., Touvron, H., Misra, I., Jegou, H., Mairal, J., Bojanowski, P., & Joulin, A. (2021). Emerging properties in self-supervised vision transformers. 2021 IEEE/CVF International Conference on Computer Vision (ICCV), 9650–9660.

Caron, M., Touvron, H., Misra, I., Jégou, H., Mairal, J., Bojanowski, P., & Joulin, A. (2021). Emerging Properties in Self-Supervised Vision Transformers. In arXiv [cs.CV]. arXiv. http://arxiv.org/abs/2104.14294

Carvalho, D. V., Pereira, E. M., & Cardoso, J. S. (2019). Machine Learning Interpretability: A Survey on Methods and Metrics. Electronics, 8(8), 832.

Castellanos-Montiel, M. J., Chaineau, M., & Durcan, T. M. (2020). The Neglected Genes of ALS: Cytoskeletal Dynamics Impact Synaptic Degeneration in ALS. Frontiers in Cellular Neuroscience, 14, 594975.

Dobin, A., Davis, C. A., Schlesinger, F., Drenkow, J., Zaleski, C., Jha, S., Batut, P., Chaisson, M., & Gingeras, T. R. (2013). STAR: ultrafast universal RNA-seq aligner. Bioinformatics, 29(1), 15–21.

Fang, M. Y., Markmiller, S., Vu, A. Q., Javaherian, A., Dowdle, W. E., Jolivet, P., Bushway, P. J., Castello, N. A., Baral, A., Chan, M. Y., Linsley, J. W., Linsley, D., Mercola, M., Finkbeiner, S., Lecuyer, E., Lewcock, J. W., & Yeo, G. W. (2019). Small-Molecule Modulation of TDP-43 Recruitment to Stress Granules Prevents Persistent TDP-43 Accumulation in ALS/FTD. Neuron, 103(5), 802–819.e11.

Fang, Z., Liu, X., & Peltz, G. (2023). GSEApy: a comprehensive package for performing gene set enrichment analysis in Python. Bioinformatics, 39(1). 10.1093/bioinformatics/btac757

Fernandopulle, M. S., Prestil, R., Grunseich, C., Wang, C., Gan, L., & Ward, M. E. (2018). Transcription Factor-Mediated Differentiation of Human iPSCs into Neurons. Current Protocols in Cell Biology / Editorial Board, Juan S. Bonifacino … [et Al.], 79(1), e51.

Fujimori, K., Ishikawa, M., Otomo, A., Atsuta, N., Nakamura, R., Akiyama, T., Hadano, S., Aoki, M., Saya, H., Sobue, G., & Okano, H. (2018). Modeling sporadic ALS in iPSC-derived motor neurons identifies a potential therapeutic agent. Nature Medicine, 24(10), 1579–1589.

Hastie, T., Friedman, J., & Tibshirani, R. (2009). The Elements of Statistical Learning. Springer New York.

Hawrot, J., Imhof, S., & Wainger, B. J. (2020). Modeling cell-autonomous motor neuron phenotypes in ALS using iPSCs. Neurobiology of Disease, 134, 104680.

Hoerl, A. E., & Kennard, R. W. (2000). Ridge Regression: Biased Estimation for Nonorthogonal Problems. Technometrics: A Journal of Statistics for the Physical, Chemical, and Engineering Sciences, 42(1), 80–86.

Hu, B.-Y., & Zhang, S.-C. (2010). Directed differentiation of neural-stem cells and subtype-specific neurons from hESCs. Methods in Molecular Biology, 636, 123–137.

Insitro, W. by. (2021, November 4). When data science goes with the flow: insitro introduces redun. Medium. https://insitro.medium.com/when-data-science-goes-with-the-flow-insitro-introduces-redun-8b06b707a14b

Klim, J. R., Williams, L. A., Limone, F., Guerra San Juan, I., Davis-Dusenbery, B. N., Mordes, D. A., Burberry, A., Steinbaugh, M. J., Gamage, K. K., Kirchner, R., Moccia, R., Cassel, S. H., Chen, K., Wainger, B. J., Woolf, C. J., & Eggan, K. (2019). ALS-implicated protein TDP-43 sustains levels of STMN2, a mediator of motor neuron growth and repair. Nature Neuroscience, 22(2), 167–179.

Landrum, M. J., Lee, J. M., Benson, M., Brown, G. R., Chao, C., Chitipiralla, S., Gu, B., Hart, J., Hoffman, D., Jang, W., Karapetyan, K., Katz, K., Liu, C., Maddipatla, Z., Malheiro, A., McDaniel, K., Ovetsky, M., Riley, G., Zhou, G.,… Maglott, D. R. (2018). ClinVar: improving access to variant interpretations and supporting evidence. Nucleic Acids Research, 46(D1), D1062–D1067.

Li, B., & Dewey, C. N. (2011). RSEM: accurate transcript quantification from RNA-Seq data with or without a reference genome. BMC Bioinformatics, 12, 323.

Lin, T.-Y., Dollár, P., Girshick, R., He, K., Hariharan, B., & Belongie, S. (2016). Feature Pyramid Networks for Object Detection. In arXiv [cs.CV]. arXiv. http://arxiv.org/abs/1612.03144

Love, M. I., Huber, W., & Anders, S. (2014). Moderated estimation of fold change and dispersion for RNA-seq data with DESeq2. Genome Biology, 15(12), 550.

Mackenzie, I. R., Rademakers, R., & Neumann, M. (2010). TDP-43 and FUS in amyotrophic lateral sclerosis and frontotemporal dementia. Lancet Neurology, 9(10), 995–1007.

Markmiller, S., Sathe, S., Server, K. L., Nguyen, T. B., Fulzele, A., Cody, N., Javaherian, A., Broski, S., Finkbeiner, S., Bennett, E. J., Lécuyer, E., & Yeo, G. W. (2021). Persistent mRNA localization defects and cell death in ALS neurons caused by transient cellular stress. Cell Reports, 36(10), 109685.

Masrori, P., & Van Damme, P. (2020). Amyotrophic lateral sclerosis: a clinical review. European Journal of Neurology: The Official Journal of the European Federation of Neurological Societies, 27(10), 1918–1929.

Melamed, Z.’ev, López-Erauskin, J., Baughn, M. W., Zhang, O., Drenner, K., Sun, Y., Freyermuth, F., McMahon, M. A., Beccari, M. S., Artates, J. W., Ohkubo, T., Rodriguez, M., Lin, N., Wu, D., Bennett, C. F., Rigo, F., Da Cruz, S., Ravits, J., Lagier-Tourenne, C., & Cleveland, D. W. (2019). Premature polyadenylation-mediated loss of stathmin-2 is a hallmark of TDP-43-dependent neurodegeneration. Nature Neuroscience, 22(2), 180–190.

Michael Ando, D., McLean, C. Y., & Berndl, M. (2017). Improving Phenotypic Measurements in High-Content Imaging Screens. In bioRxiv (p. 161422). 10.1101/161422

Mitchell, J. M., Nemesh, J., Ghosh, S., Handsaker, R. E., Mello, C. J., Meyer, D., Raghunathan, K., de Rivera, H., Tegtmeyer, M., Hawes, D., Neumann, A., Nehme, R., Eggan, K., & McCarroll, S. A. (2020). Mapping genetic effects on cellular phenotypes with “cell villages.” In bioRxiv (p. 2020.06.29.174383). 10.1101/2020.06.29.174383

Neavin, D. R., Steinmann, A. M., Farbehi, N., Chiu, H. S., Daniszewski, M. S., Arora, H., Bermudez, Y., Moutinho, C., Chan, C.-L., Bax, M., Tyebally, M., Gnanasambandapillai, V., Lam, C. E., Nguyen, U., Hernández, D., Lidgerwood, G. E., Graham, R. M., Hewitt, A. W., Pébay, A.,… Powell, J. E. (2023). A village in a dish model system for population-scale hiPSC studies. Nature Communications, 14(1), 3240.

Omole, A. E., & Fakoya, A. O. J. (2018). Ten years of progress and promise of induced pluripotent stem cells: historical origins, characteristics, mechanisms, limitations, and potential applications. PeerJ, 6, e4370.

Rayon, T., Maizels, R. J., Barrington, C., & Briscoe, J. (2021). Single-cell transcriptome profiling of the human developing spinal cord reveals a conserved genetic programme with human-specific features. Development, 148(15). 10.1242/dev.199711

Sances, S., Bruijn, L. I., Chandran, S., Eggan, K., Ho, R., Klim, J. R., Livesey, M. R., Lowry, E., Macklis, J. D., Rushton, D., Sadegh, C., Sareen, D., Wichterle, H., Zhang, S.-C., & Svendsen, C. N. (2016). Modeling ALS with motor neurons derived from human induced pluripotent stem cells. Nature Neuroscience, 19(4), 542–553.

Sareen, D., O’Rourke, J. G., Meera, P., Muhammad, A. K. M. G., Grant, S., Simpkinson, M., Bell, S., Carmona, S., Ornelas, L., Sahabian, A., Gendron, T., Petrucelli, L., Baughn, M., Ravits, J., Harms, M. B., Rigo, F., Bennett, C. F., Otis, T. S., Svendsen, C. N., & Baloh, R. H. (2013). Targeting RNA foci in iPSC-derived motor neurons from ALS patients with a C9ORF72 repeat expansion. Science Translational Medicine, 5(208), 208ra149.

Schneider, V. A., Graves-Lindsay, T., Howe, K., Bouk, N., Chen, H.-C., Kitts, P. A., Murphy, T. D., Pruitt, K. D., Thibaud-Nissen, F., Albracht, D., Fulton, R. S., Kremitzki, M., Magrini, V., Markovic, C., McGrath, S., Steinberg, K. M., Auger, K., Chow, W., Collins, J.,… Church, D. M. (2017). Evaluation of GRCh38 and de novo haploid genome assemblies demonstrates the enduring quality of the reference assembly. Genome Research, 27(5), 849–864.

Shao, W., Todd, T. W., Wu, Y., Jones, C. Y., Tong, J., Jansen-West, K., Daughrity, L. M., Park, J., Koike, Y., Kurti, A., Yue, M., Castanedes-Casey, M., Del Rosso, G., Dunmore, J. A., Zanetti Alepuz, D., Oskarsson, B., Dickson, D. W., Cook, C. N., Prudencio, M.,… Petrucelli, L. (2022). Two FTD-ALS genes converge on the endosomal pathway to induce TDP-43 pathology and degeneration. Science, 378(6615), 94–99.

Sivanandan, S., Leitmann, B., Lubeck, E., Sultan, M. M., Stanitsas, P., Ranu, N., Ewer, A., Mancuso, J. E., Phillips, Z. F., Kim, A., Bisognano, J. W., Cesarek, J., Ruggiu, F., Feldman, D., Koller, D., Sharon, E., Kaykas, A., Salick, M. R., & Chu, C. (2023). A Pooled Cell Painting CRISPR Screening Platform Enables de novo Inference of Gene Function by Self-supervised Deep Learning. In bioRxiv (p. 2023.08.13.553051). 10.1101/2023.08.13.553051

Streit, L., Kuhn, T., Vomhof, T., Bopp, V., Ludolph, A. C., Weishaupt, J. H., Gebhardt, J. C. M., Michaelis, J., & Danzer, K. M. (2022). Stress induced TDP-43 mobility loss independent of stress granules. Nature Communications, 13(1), 5480.

Stringer, C., Wang, T., Michaelos, M., & Pachitariu, M. (2021). Cellpose: a generalist algorithm for cellular segmentation. Nature Methods, 18(1), 100–106.

Tange, O. (2023). GNU Parallel 20230222 (’Gaziantep’) released. Zenodo. 10.5281/ZENODO.7668337

Thams, S., Lowry, E. R., Larraufie, M.-H., Spiller, K. J., Li, H., Williams, D. J., Hoang, P., Jiang, E., Williams, L. A., Sandoe, J., Eggan, K., Lieberam, I., Kanning, K. C., Stockwell, B. R., Henderson, C. E., & Wichterle, H. (2019). A Stem Cell-Based Screening Platform Identifies Compounds that Desensitize Motor Neurons to Endoplasmic Reticulum Stress. Molecular Therapy: The Journal of the American Society of Gene Therapy, 27(1), 87–101.

Tibshirani, R. (1996). Regression shrinkage and selection via the lasso. Journal of the Royal Statistical Society. Series B, Statistical Methodology, 58(1), 267–288.

van Eersel, J., Ke, Y. D., Gladbach, A., Bi, M., Götz, J., Kril, J. J., & Ittner, L. M. (2011). Cytoplasmic accumulation and aggregation of TDP-43 upon proteasome inhibition in cultured neurons. PloS One, 6(7), e22850.

Verzat, C., Harley, J., Patani, R., & Luisier, R. (2022). Image-based deep learning reveals the responses of human motor neurons to stress and VCP-related ALS. Neuropathology and Applied Neurobiology, 48(2), e12770.

Weishaupt, J. H., Hyman, T., & Dikic, I. (2016). Common Molecular Pathways in Amyotrophic Lateral Sclerosis and Frontotemporal Dementia. Trends in Molecular Medicine, 22(9), 769–783.

Workman, M. J., Lim, R. G., Wu, J., Frank, A., Ornelas, L., Panther, L., Galvez, E., Perez, D., Meepe, I., Lei, S., Valencia, V., Gomez, E., Liu, C., Moran, R., Pinedo, L., Tsitkov, S., Ho, R., Kaye, J. A., Answer ALS Consortium,… Svendsen, C. N. (2023). Large-scale differentiation of iPSC-derived motor neurons from ALS and control subjects. Neuron, 111(8), 1191–1204.e5.

Wu, J., Chen, S., Liu, H., Zhang, Z., Ni, Z., Chen, J., Yang, Z., Nie, Y., & Fan, D. (2018). Tunicamycin specifically aggravates ER stress and overcomes chemoresistance in multidrug-resistant gastric cancer cells by inhibiting N-glycosylation. Journal of Experimental & Clinical Cancer Research: CR, 37(1), 272.

